# A generic cell surface ligand system for studying cell-cell recognition

**DOI:** 10.1101/546846

**Authors:** Eleanor M Denham, Michael I Barton, Susannah M Black, Marcus J Bridge, Ben de Wet, Rachel L Paterson, P. Anton van der Merwe, Jesse Goyette

**Affiliations:** Sir William Dunn School of Pathology, University of Oxford, Oxford, UK; School of Medical Sciences, University of New South Wales, Sydney, Australia

## Abstract

Dose-response experiments are a mainstay of receptor biology studies and can reveal valuable insights into receptor function. Such studies of receptors that bind cell surface ligands are currently limited by the difficulty in manipulating the surface density of ligands at a cell-cell interface. Here we describe a generic cell surface ligand system that allows precise manipulation of cell surface ligand densities over several orders of magnitude. We validate the system for a range of immunoreceptors, including the T cell receptor (TCR), and show that this generic ligand stimulates via the TCR at a similar surface density as its native ligand. This system allows the effect of surface density, valency, dimensions, and affinity of the ligand to be manipulated. It can be readily extended to other receptor-cell surface ligand interactions, and will facilitate investigation into the activation of, and signal integration between, cell surface receptors.

## Introduction

Many cellular receptors are activated by ligands presented on other cell surfaces. To study these receptors in depth, cell lines expressing the appropriate physiological ligand are required. Conducting dose-response experiments on these receptors is challenging as controlling ligand density on the surface of cells is difficult. Currently this is limited to sorting cells into populations with varying expression levels, using inducible expression systems, or the use of blocking antibodies. An alternative to physiological ligands that is commonly used is antibodies specific for a receptor or recombinant ligands presented on artificial surfaces, such as plastic or glass-supported lipid bilayers. However, this can be a poor mimic for the complexity and biophysical characteristics of cell surfaces.

One widely studied group of receptors are non-catalytic tyrosine-phosphorylated receptors (NTRs), also called immunoreceptors, which are the largest group of receptors expressed on leukocytes^1^. These receptors play a major role in the recognition of infected or cancer cells. Since some NTRs, such as the T cell receptor (TCR), cytotoxic T-lymphocyte–associated antigen-4 (CTLA-4) and programmed death-1 (PD-1), have been successfully manipulated/targeted for therapeutic purposes^2,3^, the remaining NTRs are currently under intense investigation for the development of immunotherapies (examples include^4–7^).

The mechanism of signal transduction, or triggering, has been studied in depth for some NTRs (notably the TCR) but remains controversial^1,8^. Due to the widespread importance of NTR function in immune regulation, and huge interest in their activity, elucidating this mechanism is critical to both our understanding and ability to modulate receptor activity where required in clinical settings. While NTRs have conserved signalling modules, their extracellular regions are rapidly evolving, hugely diverse, and bind a structurally-diverse range of ligands^1^. This diversity, and the fact that their ligands are often not known, has hampered a systematic investigation of NTRs.

These limitations motivated us to develop new tools for investigating receptors that have cell surface ligands. We present a novel generic ligand system whereby a single ligand can engage any receptor engineered to have an accessible tag. We show that ligand density can be varied easily and precisely quantified and we show that the generic ligand can activate a number of representative NTRs in a clear dose-dependent manner. For one receptor, the TCR, we compare the response to generic versus physiological ligand: peptide presented in major histocompatibility complexes (pMHC).

Finally, we describe and briefly show how this generic ligand can be manipulated to alter other biochemical and biophysical properties of receptor-ligand interactions such as dimensions, affinity and valency. These modifications to a single ligand will apply to any interaction involving the ligand and a tagged receptor and thus permit high-throughput, systematic analyses of multiple receptors.

## Results

### Design and development of a generic ligand system

We aimed to design a system requiring minimal manipulation of the receptor, in order to preserve its structure and interactions. In addition, we sought to create a cell surface-expressed ligand that would engage receptors with an affinity comparable to physiological NTR-ligand interactions and which, when bound to receptor, would preserve the cell-cell intermembrane distance.

We developed a generic ligand based on the interaction between the peptide Twin-Strep-tag, which has two Strep-tag II motifs, and Strep-Tactin, a variant of streptavidin (outlined in Fig. 1) ^9–11^.

**Figure 1:**
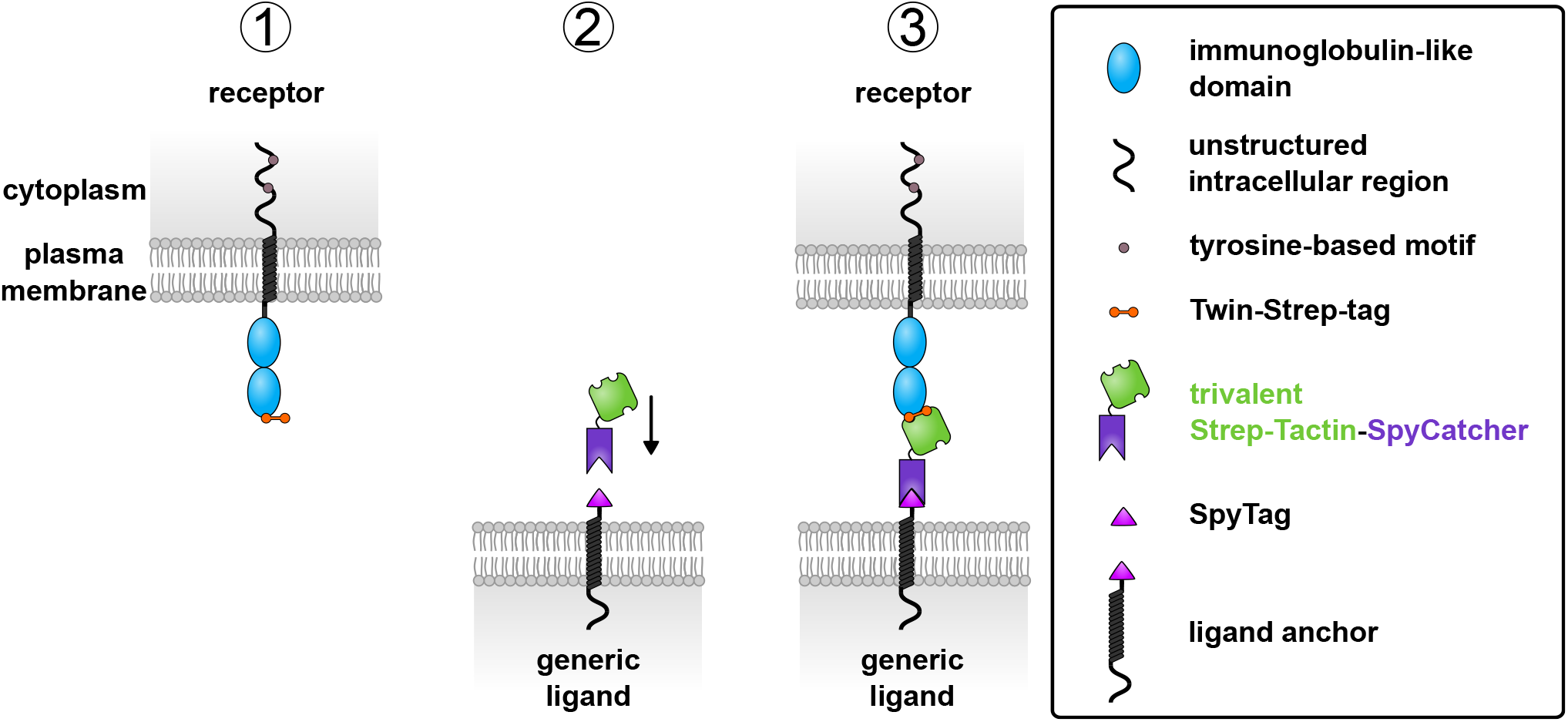
Design of a generic ligand system. (1) Each receptor is constructed with a Twin-Strep-tag at the extracellular terminus and expressed in cell lines. (2) The generic ligand is made of two components: a cell surface-expressed ligand anchor with N-terminal SpyTag, and soluble trivalent Strep-Tactin-SpyCatcher protein. Trivalent Strep-Tactin-SpyCatcher is added to cells expressing the generic ligand anchor. When Spy-Catcher and SpyTag interact, a spontaneous covalent isopeptide bond forms between them creating the complete generic ligand. (3) The three binding sites of trivalent Strep-Tactin-SpyCatcher are available for ligation by the Twin-Strep-tagged receptor. Two binding sites are required for the full engagement of Twin-Strep-tag.

The NTR of interest, expressed on a “receptor” cell, is genetically engineered to add the Twin-Strep-tag peptide to the extracellular N-terminus where it is accessible for engagement by the generic ligand presented on another “ligand” cell (Fig. 1). This ligand is made up of two components: a cell surface ligand anchor and a soluble fusion protein that spontaneously forms a covalent bond with the anchor. The ligand anchor comprises the transmembrane and cytoplasmic portions of mouse CD80 fused to an N-terminal SpyTag peptide, forming one half of the covalent bond-forming split-protein pair SpyTag/SpyCatcher^12–14^. The soluble fusion protein comprises trivalent Strep-Tactin, which has three binding sites for Strep-tag II motifs, fused to SpyCatcher (trivalent Strep-Tactin-SpyCatcher) ^9,10^. When soluble trivalent Strep-Tactin-SpyCatcher is incubated with cells expressing the ligand anchor, SpyTag and SpyCatcher spontaneously form a covalent bond. This yields a cell surface ligand able to bind a Twin-Strep-tagged receptor (Fig. 1).

Based on the available structures of Strep-Tactin and SpyTag/SpyCatcher, and estimates of linker lengths, we predict the complete generic ligand to have an extracellular length similar to 2-3 immunoglobulin-like domains when extended (PDB accession numbers: 1KL3, 4MLI) ^15,16^. This is comparable to physiological NTR ligands (discussed in Dushek et al. ^1^).

### Preparation of trivalent Strep-Tactin-SpyCatcher

To prepare trivalent Strep-Tactin-SpyCatcher tetramers, we employed a method previously used to generate streptavidin tetramers of defined valency^17,18^. This uses a mutated streptavidin subunit that has negligible biotin-binding activity, termed “dead” streptavidin. Biotin and Strep-tag II occupy the same surface pocket of streptavidin and so we assumed that the dead streptavidin subunit is also unable to bind Strep-tag II^19^. Strep-Tactin subunits were refolded from bacterial inclusion bodies with subunits of dead streptavidin fused at its C-terminus to SpyCatcher (dead streptavidin-SpyCatcher) in a 3:1 molar ratio (Supplementary Fig. 1).

Dead streptavidin-SpyCatcher contains a polyaspartate insertion allowing purification of the trivalent Strep-Tactin-SpyCatcher tetramer from other possible configurations using anion exchange chromatography (Supplementary Fig. 1).

We analysed a sample from the first eluted peak, predicted to be that of trivalent Strep-Tactin-SpyCatcher, using SDS-PAGE. Upon boiling, the tetramer is reduced to individual monomers of Strep-Tactin and dead streptavidin-SpyCatcher allowing visualisation of the relative proportion of the subunits (Supplementary Fig. 1). For comparison we analysed a sample from a later peak, which we predict to contain monovalent Strep-Tactin (1 x Strep-Tactin, 3 x dead streptavidin-SpyCatcher) based on order of elution. The ratio of Strep-Tactin to dead streptavidin-SpyCatcher subunits was higher than expected (4.7:1 for trivalent Strep-Tactin-SpyCatcher) but was consistent with a 3:1 ratio when compared to the subunit ratio of the monovalent protein (Supplementary Fig. 1).

To confirm that the purified protein was the desired tetramer we used a biotin-4-fluorescein fluorescence quenching assay previously used to estimate the number of biotin-binding sites per streptavidin tetramer (Supplementary Fig. 1) ^18^. When bound to Strep-Tactin the fluorescence of biotin-4-fluorescein is quenched. As the concentration of biotin-4-fluorescein added to trivalent Strep-Tactin-SpyCatcher increases above binding site saturation, there is an increasing amount of free (non-quenched) biotin-4-fluorescein in solution. This is visualised as a sharp increase in fluorescence with the inflection point indicating saturation. Titration of biotin-4-fluorescein yielded an estimated number of 4.2 binding sites per trivalent Strep-Tactin-SpyCatcher (Supplementary Fig. 1). While higher than the expected value of 3, it is three times the estimated binding site number calculated for the predicted monovalent Strep-Tactin peak (1.4). We assume there are the same number of available Strep-tag II binding sites per tetramer.

### Characterisation of the generic ligand

#### Affinity between Twin-Strep-tag and trivalent Strep-Tactin-SpyCatcher

We used surface plasmon resonance and a Twin-Strep-tag-monomeric Teal Fluorescent Protein (mTFP) fusion protein to analyse Twin-Strep-tag binding to trivalent Strep-Tactin-SpyCatcher at 37°C. Using data from seven independent experiments (n = 11), the K_D_ was measured as 6.8 µM (Fig. 2a). This is comparable to the affinities reported for physiological NTR-ligand interactions we wish to replicate^20–22^.

**Figure 2:**
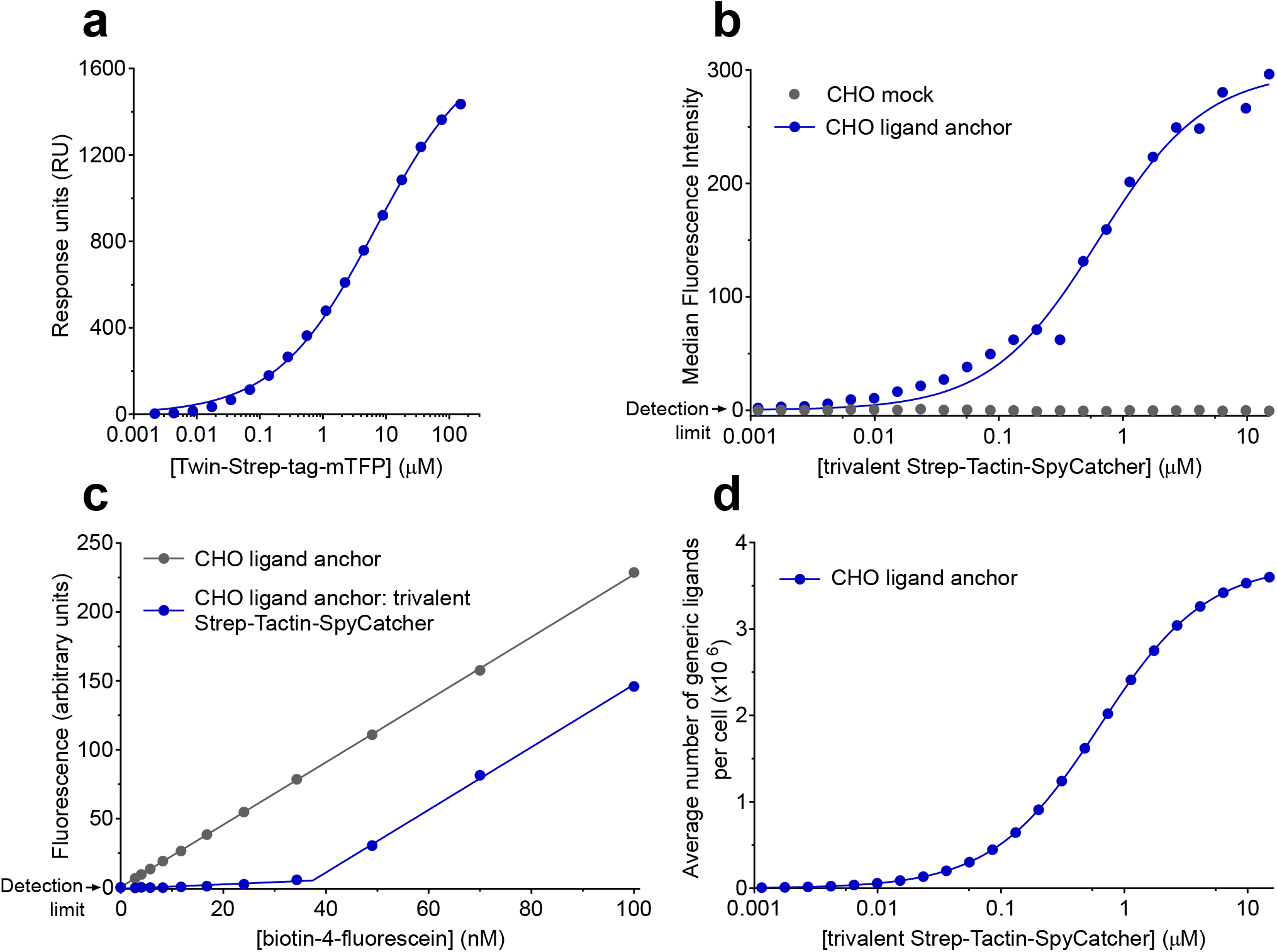
Characterisation of Twin-Strep-tag: trivalent Strep-Tactin-SpyCatcher interaction and quantifying the number of generic ligands per cell. (**a**) Representative equilibrium binding measured by surface plasmon resonance of Twin-Strep-tag-mTFP injected over immobilised trivalent Strep-Tactin-Spycatcher at 37 °C is shown. The K_D_ (standard error of the mean, SEM) for the collated data from seven independent experiments (n = 11) is 6.8 µM (0.62 µM) and the mean Hill slope (SEM) is 0.46 (0.03) to 2 significant figures (s.f.). (**b**) A relative indication of the level of generic ligand per cell as a function of trivalent Strep-Tactin-SpyCatcher concentration added to cells. Median fluorescence intensity values extracted from flow cytometry analyses of cells incubated with ATTO 647 biotin are shown. (**c**) CHO ligand anchor cells pre-incubated with trivalent Strep-Tactin-SpyCatcher or buffer alone were incubated with a titration of biotin-4-fluorescein in a fluorescence quenching assay. The inflection point is used to calculate average absolute number of generic ligands per cell. (**d**) The saturating concentration of biotin-4-fluorescein was extracted from c and converted into number of generic ligands per cell. This was substituted as the maximum into the fitted curve in b to interpolate the average number of generic ligands per cell as a function of trivalent Strep-Tactin-SpyCatcher concentration added to cells.

#### Characterising the optimal conditions for ligand anchor:trivalent Strep-Tactin-SpyCatcher binding

A stable, high-expressing ligand anchor cell line was established and maintained under selection (Supplementary Fig. 2). Chinese Hamster Ovary (CHO) cells were used to avoid any confounding receptor-ligand interactions that might occur if both receptor and ligand cells were human.

We explored the optimal conditions for covalent coupling between the cell surface-presented ligand anchor and soluble trivalent Strep-Tactin-SpyCatcher. In line with the findings of Zakeri and colleagues, we found that coupling between CHO ligand anchor cells and trivalent Strep-Tactin-SpyCatcher was most efficient in buffer at pH 5-6 (Supplementary Fig. 2) ^12^. The widest range of surface densities was achieved with a 10 minute incubation of trivalent Strep-Tactin-SpyCatcher at a wide range of concentrations (Supplementary Fig. 2).

Using western blotting on boiled cell lysates fractionated by SDS-PAGE we visualised the ligand anchor by probing for the N-terminal HA tag (Supplementary Fig. 2). Addition of trivalent Strep-Tactin-SpyCatcher to the cells led to a substantial increase in the molecular weight of a significant proportion of ligand anchor consistent with covalent coupling to the soluble fusion protein. Probing with anti-streptavidin antibody confirmed this (Supplementary Fig. 2). A time course showed that a significant proportion of generic ligand remains at the cell surface for many hours post-reconstitution (Supplementary Fig. 2). Receptor stimulation assays are commonly conducted over this time frame so the ligand is able to provide a strong stimulus for this duration.

### Measuring the number of generic ligands per cell

A major strength of the generic ligand system is the ligand dose can be varied easily by titrating the concentration of soluble trivalent Strep-Tactin-SpyCatcher added to cells. To enable us to determine the ligand surface density required for activation we developed a method to measure the number of generic ligands per cell. This method uses two assays. The first, whereby CHO generic ligand cells are incubated with ATTO 647 biotin, gives an indication of how the relative number of generic ligand sites varies with Strep-Tactin-SpyCatcher concentration (Fig. 2b).

The second assay measures the average maximum number of generic ligands per cell in a population of cells saturated with trivalent Strep-Tactin-SpyCatcher. This uses the same biotin-4-fluorescein fluorescence quenching assay shown in Supplementary Fig. 1. Assuming complete binding, the biotin-4-fluorescein concentration that saturates the cells is indicated by the inflection point of the graph (Fig. 2c). This saturating concentration for a known number of cells in a defined volume can then be converted into an average number of generic ligands per cell (see Methods). By combining the absolute number of generic ligands per cell at saturation (Fig. 2c) and the relative ligand levels across a range of soluble trivalent Strep-Tactin-SpyCatcher concentrations (Fig. 2b), the average number of generic ligands per cell for a given soluble trivalent Strep-Tactin-SpyCatcher concentration can be estimated (Fig. 2d). A maximum of 3 million generic ligands per cell can be consistently achieved and the ligand dose can therefore easily be varied over several orders of magnitude.

### Representative activating NTRs can be stimulated by generic ligand

Representative activating human NTRs from various receptor families were genetically modified to have an N-terminal Twin-Strep-tag (Fig. 3a).

**Figure 3:**
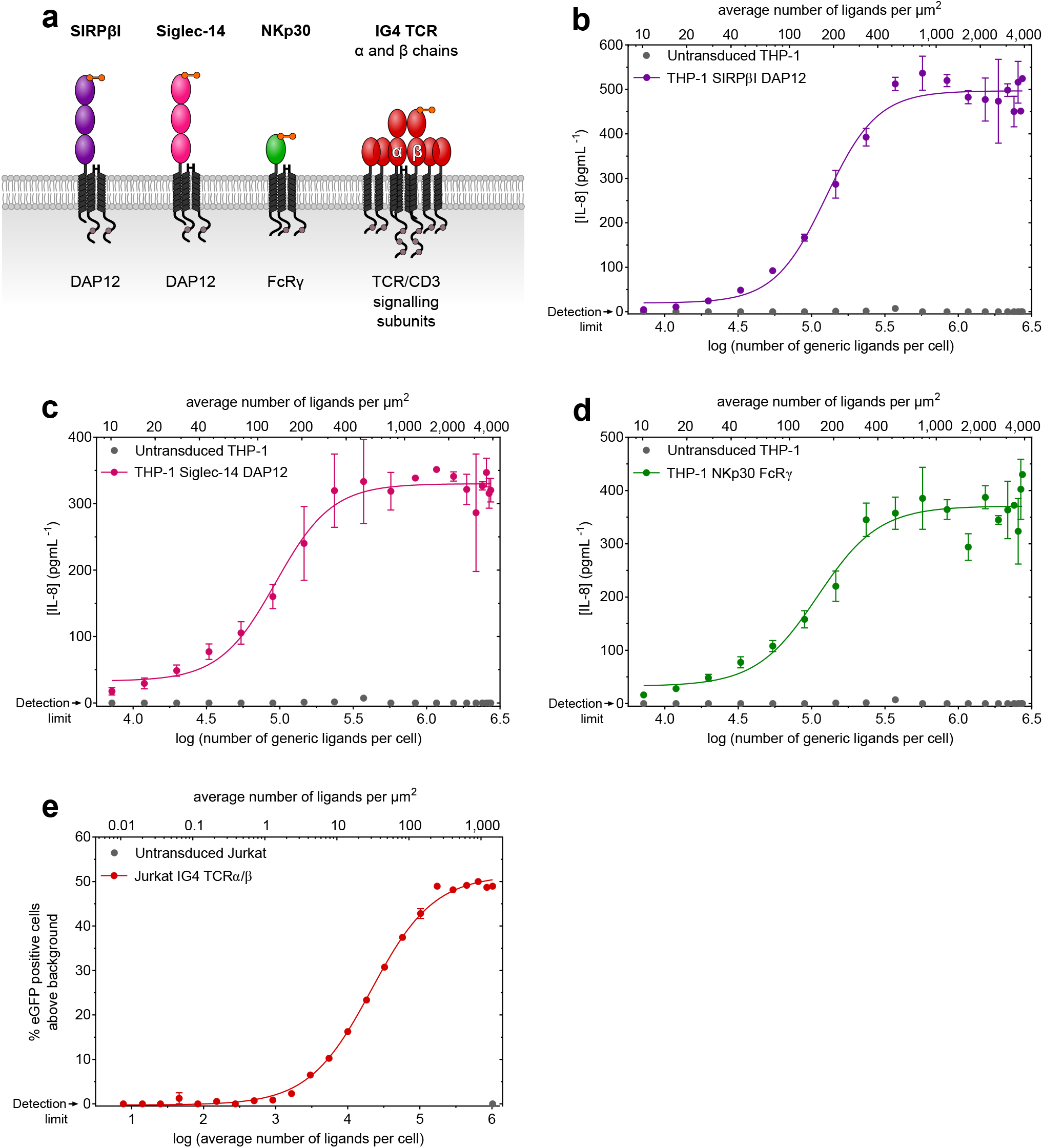
Twin-Strep-tagged receptors can be stimulated by generic ligand-presenting cells. (**a**) Cartoon depictions of four representative NTRs with Twin-Strep-tags, and adaptor proteins, are shown. In the case of IG4 TCR, the β chain has an N-terminal Twin-Strep-tag. The extracellular region of each receptor contains one or more immunoglobulin-like domains (see legend in Fig. 1). Response of Twin-Strep-tagged SIRPβI (**b**), Siglec-14 (**c**), or NKp30 (**d**) expressing THP-1 cells to generic ligand presented on CHO cells. Receptor response is measured by IL-8 secretion. (**e**) Response of Jurkat IG4 TCRα/β-Twin-Strep-tag cells to generic ligand presented on CHO cells. Receptor response, indicated by eGFP expression under the control of the NFκB promoter, is shown as percentage of cells positive for eGFP above background. Error bars indicate the range (n = 2) and data are representative of three independent experiments with the collated data shown in table 1. Within each stimulation, a sample of CHO cells were taken and used to measure the number of generic ligands per cell as in Fig. 2. Ligand density was calculated from these numbers using an estimated CHO cell surface area of 700 µm^2^ (see methods) ^47,48^. DAP12 - DNAX-activating protein of 12 kDa.

Signal regulatory protein βI (SIRPβI, CD172b) and Siglec-14 are expressed in a number of leukocytes including monocytes and macrophages^23,24^. Siglec-14 binds to sialic acid presented by numerous bacteria to induce responses including cytokine secretion^24^. There is evidence that SIRPβI contributes to neutrophil transepithelial migration and macrophage phagocytosis, but a ligand has yet to be identified^23^. A generic ligand is therefore very useful for the study of SIRPβI. As a member of the natural killer cell cytotoxicity family, NKp30 (CD337) is expressed in natural killer (NK) cells and binds both pathogen and cellular ligands to mediate NK cell cytotoxicity^25^.

SIRPβI, Siglec-14, and NKp30 receptors each associate with adaptor proteins that contain cytoplasmic phosphorylatable tyrosine residues that mediate immune signalling^23–25^. Therefore, to establish stable cell lines, THP-1 cells were co-transduced with lentiviruses encoding the tagged receptor and appropriate adaptor (Supplementary Fig. 3).

The αβT cell receptor (TCR) complex consists of TCR α and β chains associated with a TCRζ homodimer, and CD3δϵ and CD3γϵ heterodimers. The IG4 TCR α chain and Twin-Strep-tagged β chain were transduced into a Jurkat nuclear factor kappa-light-chain-enhancer of activated B cells (NFκB) reporter cell line in which the production of enhanced Green Fluorescent Protein (eGFP) is under the control of the NFκB promoter (Supplementary Fig. 3) ^26,27^. Any IG4 TCR α and β chains expressed at the cell surface are presumed to be associated with endogenous TCR/CD3 signalling subunits.

All four tagged receptors were activated by generic ligand cells with a clear dose-dependent response which increased with the number of generic ligands per cell (Fig. 3b-e). This response was specific as shown by the absence of IL-8 secretion or NFκB-driven eGFP expression in samples containing negative control receptor cells (Fig. 3b-e) or ligand cells (data not shown). The EC_50_ and hill slope values of the receptor responses from three independent experiments are shown in Table 1. Thus the generic ligand can bind to and trigger several different receptors bearing an N-terminal Twin-Strep-tag.

**Table 1:** Collated EC_50_ and hill slope values from receptor stimulation experiments. The mean EC_50_ and hill slope values along with standard errors of the mean of THP-1 SIRPβI DAP12, THP-1 Siglec-14 DAP12, or THP-1 NKp30 FcRγ cell response to generic ligand from three independent experiments are shown. The mean EC_50_ and hill slope values along with standard errors of the mean of Jurkat IG4 TCRα/β cell response to generic ligand or 9V-HLA-A*02 from three independent experiments are also shown. EC_50_ values are given as both number of generic ligands or 9V-HLA-A*02 per cell and ligand density to 2 significant figures (s.f.). Hill slope values are also given to 2 s.f.

|  | mean EC<sub>50</sub> ± standard error of the mean (ligands per cell) | mean EC<sub>50</sub> ± standard error of the mean (ligands per µm<sup>2</sup>) | mean hill slope ± standard error of the mean |
| --- | --- | --- | --- |
| **Response to generic ligand** |  |  |  |
| THP-1 SIRPβI DAP12 | 260,000 ± 110,000 | 360 ± 160 | 1.8 ± 0.50 |
| THP-1 Siglec-14 DAP12 | 190,000 ± 99,000 | 260 ± 140 | 1.4 ± 0.45 |
| THP-1 NKp30 FcRγ | 130,000 ± 24,000 | 190 ± 33 | 1.5 ± 0.33 |
| Jurkat IG4 TCRα/β | 17,000 ± 3,100 | 23 ± 4.4 | 0.97 ± 0.070 |
| **Response to 9V-HLA-A\*02** |  |  |  |
| Jurkat IG4 TCRα/β | 7,100 ± 3,600 | 10 ± 5.1 | 1.0 ± 0.054 |

### Twin-Strep-tagged T cell receptor responds to generic ligand and native ligand with a similar sensitivity

In order to validate this approach we compared the response of the TCR to generic ligand with its response to native ligand, peptide-MHC.

IG4 TCR and its cognate ligand NY-ESO-1_157-165_ 9V peptide variant (SLLMWITQV) presented in complex with HLA-A*02 is a well characterised receptor-ligand pair^28–30^. The dissociation constant of IG4 TCR binding to 9V-HLA-A*02 (K_D_ = 6-7 µM) is comparable to the K_D_ we have measured for Twin-Strep-tag and trivalent Strep-Tactin-SpyCatcher (Fig. 2a) ^29,30^.

To compare the TCR response to either generic or native ligand, the Jurkat NFκB reporter cell line transduced with IG4 TCR α and Twin-Strep-tagged β chains (Supplementary Fig. 3) was presented to CHO cells that express both HLA-A*02 in the form of a single chain dimer and the generic ligand anchor (Fig. 4a, Supplementary Fig. 3). These CHO cells were either pre-incubated with trivalent Strep-Tactin-SpyCatcher or loaded with 9V peptide. To allow a direct comparison between generic and pMHC ligand the Jurkat cells were not transduced with the co-receptor CD8, since it binds MHC but not generic ligand.

**Figure 4:**
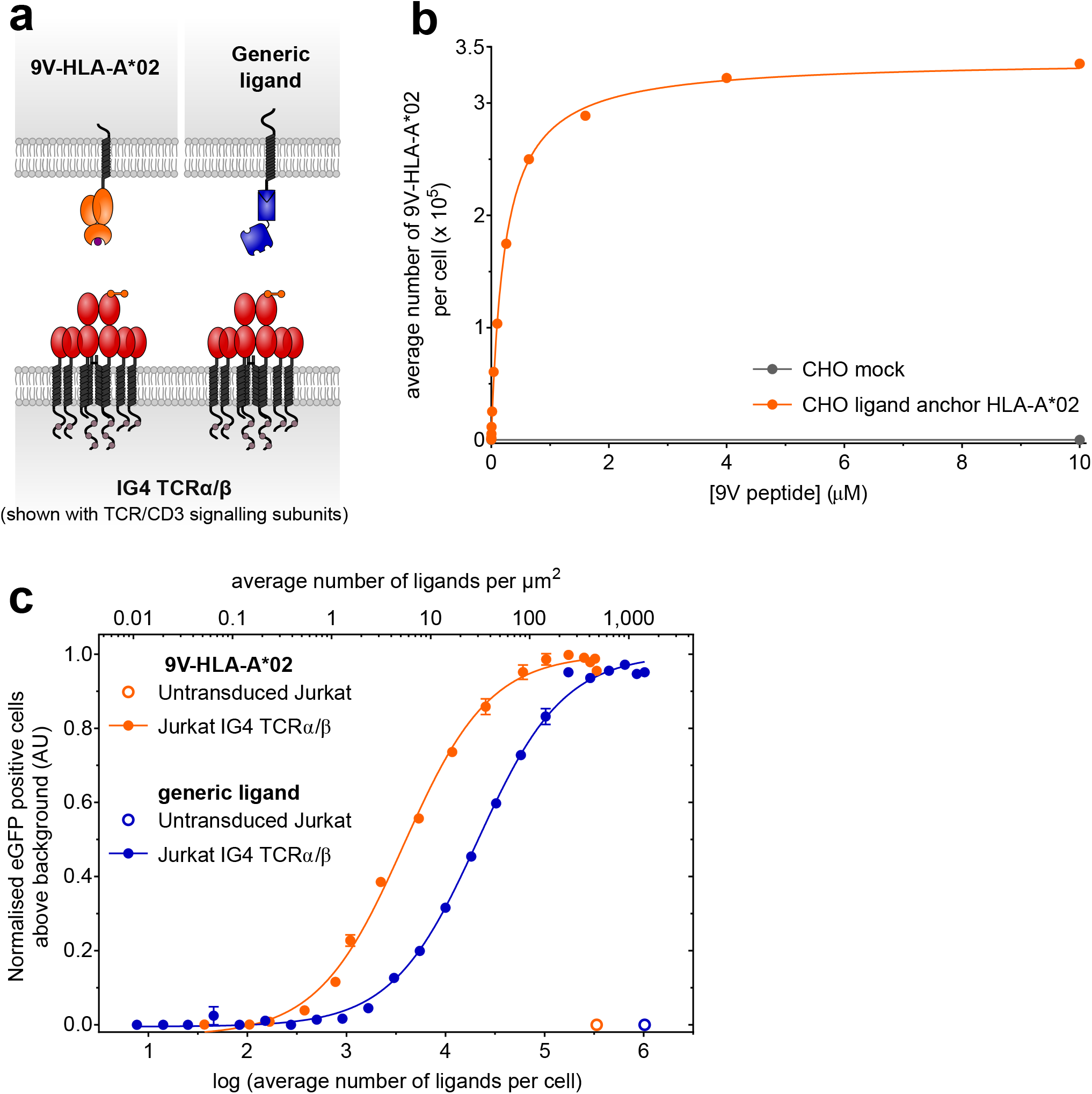
IG4 TCR responds to both generic and physiological ligand with a similar sensitivity. (**a**) A cartoon depicting IG4 TCR with β chain N-terminal Twin-Strep-tag presented to either 9V-HLA-A*02 or generic ligand on CHO cells. (**b**) The average number of 9V-HLA-A*02 per cell as a function of peptide concentration added was measured using soluble fluorescent IG4 high affinity TCR and fluorescence quantitation beads. Interpolated numbers of 9V-HLA-A*02 are shown. (**c**) Response of Jurkat IG4 TCRα/β cells to either generic ligand or 9V-HLA-A*02 presented on CHO cells. Receptor response, as indicated by eGFP expression under the control of the NFκB promoter, is shown normalised to the maximal receptor response to either 9V-HLA-A*02 or generic ligand. Error bars indicate the range (n = 2) and data are representative of three independent experiments with the collated data shown in table 1.

In order to quantitatively compare the TCR response to generic ligand or 9V-HLA-A*02, we measured the number of 9V-HLA-A*02 molecules on the CHO ligand cells. For this, we used a soluble affinity-enhanced (c58/c61) form of the IG4 TCR that binds to 9V-HLA-A*02 with a much higher affinity than the wild-type TCR (K_D_ = 71 pM) ^31,32^. This soluble IG4 high affinity TCR was labelled with Alexa Fluor 647 (IG4hi TCR AF647) and used in combination with fluorescence quantitation beads to interpolate the average number of 9V-HLA-A*02 per cell (Fig. 4b, Supplementary Fig. 4). Incubating CHO ligand anchor HLA-A*02 cells with varying concentrations of 9V peptide yielded a large dynamic range of ligand number per cell (Fig. 4b). Therefore, we were able to present Jurkat IG4 TCRα/β-Twin-Strep-tag cells with CHO ligand anchor HLA-A*02 cells presenting either 9V-HLA-A*02 or generic ligand at similar densities.

Jurkat IG4 TCRα/β-Twin-Strep-tag cells responded to both 9V-HLA-A*02 and generic ligand in a dose-dependent, specific manner, visualised using the NFκB reporter (Fig. 4c). The EC_50_ and hill slope values from three independent experiments are shown in Table 1. There is a 2-fold difference in the average sensitivity between the receptor response to 9V-HLA-A*02 versus generic ligand. We believe the variability of 9V-HLA-A*02 EC_50_ values arises from inter-experiment variability in estimating the number of 9V-HLA-A*02 molecules per cell.

Based on the structure of soluble IG4 TCRα/β bound to cognate pMHC we predict the presence of Twin-Strep-tag should not interfere with TCR-pMHC binding (PDB accession number: 2BNR) ^29^. We used soluble pMHC class I tetramer staining to confirm this. Because tetramer staining of Jurkat IG4 TCRα/β cells in the absence of CD8 co-receptor expression was very poor (data not shown) we used cells that express CD8. Jurkat cells expressing either non-tagged, Strep-tag II-tagged or Twin-Strep-tagged IG4 TCRα/β, and CD8α/β, were matched for TCRβ chain and CD8α expression (Supplementary Fig. 4). These cells showed comparable cognate pMHC tetramer staining (Supplementary Fig. 4) suggesting the presence of Strep-tag II does not significantly interfere with pMHC binding.

In summary, we have demonstrated that the generic ligand can elicit a receptor response comparable to physiological ligand, reinforcing its usefulness for studying receptor activation.

### Manipulating the generic ligand system

Our generic ligand system could be adapted to allow manipulation of ligand properties other than surface density, such as length, affinity, and valency, as illustrated in Fig. 5a-c. For example, the receptor can be tagged with an N-terminal Strep-tag II and presented to monovalent Strep-Tactin-SpyCatcherΔ on cells, yielding a receptor-ligand pair with a 6-fold higher dissociation constant than the Twin-Strep-tag-trivalent Strep-Tactin pair, (K_D_ = 43 µM from 3 independent experiments, n = 6) (highlighted in Fig. 5b, Supplementary Fig. 5). Preparation of monovalent Strep-Tactin-SpyCatcherΔ and quantitation of monovalent Strep-Tactin-SpyCatcherΔ ligand numbers per cell are performed in the same manner as for the higher affinity system (Supplementary Fig. 5-6). Siglec-14 with N-terminal Strep-tag II and expressed with its adaptor in THP-1 cells was able to respond to monovalent Strep-Tactin-SpyCatcherΔ presented on CHO cells (Fig. 5d). As we would predict, the EC_50_ of this response to a lower affinity interaction was higher, by approximately six-fold, compared to that measured for Siglec-14 with Twin-Strep-tag and trivalent Strep-Tactin-SpyCatcher (mean EC_50_ values of 1,100,000 and 190,000 generic ligands per cell respectively).

**Figure 5:**
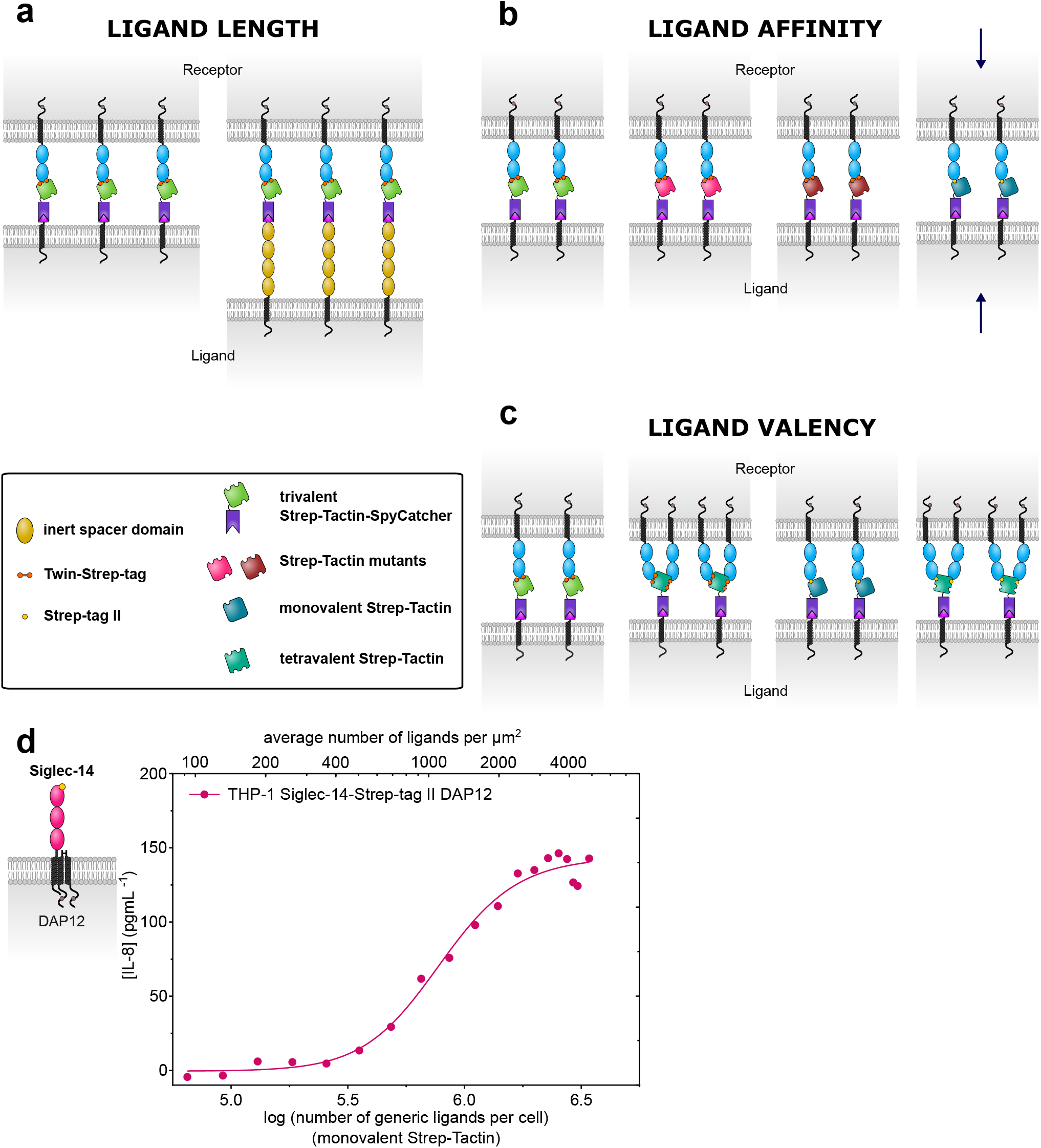
Manipulations of the generic ligand system. (**a**) The extracellular region of the generic ligand can be elongated by insertion of inert spacer domains. Receptor responses to cells expressing this elongated ligand can be compared to responses of receptors ligated with the shorter ligand. (**b**) Mutated forms of Strep-Tactin/streptavidin or receptors with a single Strep-tag II can be substituted into the receptor-generic ligand interaction to investigate how changing binding affinity affects receptor activation and signalling. Strep-Tactin variants are shown in different colours. One such affinity variant (indicated by arrows) involves presenting Strep-tag II-tagged receptors to monovalent Strep-Tactin. (**c**) Strep-Tactin-SpyCatcher (or streptavidin-SpyCatcher) tetramers with varying numbers of Strep-tag II binding sites can be coupled to CHO ligand anchor cells to examine the effect of varying the valency of the generic ligand on NTR triggering. (**d**) Response of THP-1 Siglec-14-Strep-tag II DAP12 cells to monovalent Strep-Tactin generic ligand presented on CHO cells. Receptor response is measured by IL-8 secretion. Data are from one of three independent experiments that together have a mean EC_50_ ± standard error of the mean of 1,100,000 ± 120,000 generic ligands per cell.

## Discussion

We describe a novel generic ligand system that facilitates the investigation of receptors that bind cell surface ligands. This ligand is stably presented on the surface of CHO cells, and the ligand density can be easily titrated over several orders of magnitude. We believe this fine control over ligand dose within a cell environment is a clear advantage over current methods with physiological ligand.

We show that four representative NTRs can be stimulated by ligation of N-terminal Twin-Strep-tag. These results show that specific engagement of native ligand is not required for receptor triggering. This is in agreement with other work that shows these NTRs, among others, can be activated by antibodies to the receptor ectodomains coated on plastic (examples include^8,23,25^).

Twin-Strep-tagged SIRPβI, Siglec-14 and NKp30 respond to Strep-Tactin generic ligand with a similar sensitivity. The mean EC_50_ values of the receptor responses from three independent experiments were 130,000-260,000 ligands per cell (Table 1). In contrast, the IG4 TCR responds with a much lower mean EC_50_ of 17,000 ligands per cell, indicating that TCR triggering required much lower surface densities of generic ligand than other NTRs (Table 1). Furthermore, the ligand densities of our generic ligand required to elicit responses were comparable to the required ligand densities of native ligands reported by others^33–37^.

A number of manipulations can be made to the generic ligand system to investigate the basic requirements and optimal conditions for NTR activation, as outlined in Fig. 5a-c. Increasing the extracellular dimensions of the generic ligand by inserting inert spacer domains into the anchor would enable testing whether ligand length affects recognition, as predicted by the kinetic-segregation model of NTR triggering (Fig. 5a) ^1,38^. Until now these experiments have been painstaking as they require expressing different length forms of each native ligand at matched levels, and titration of ligand density is not possible^1,39^.

We also illustrate how the receptor-generic ligand affinity can be altered through using alternative SpyCatcher fusion proteins and/or variants of the Strep-tag II/Twin-Strep-tag system (Fig. 5b,d, Supplementary Fig. 5-6). Thus the sensitivity of receptors to affinity changes, and whether there is an optimal ligand binding affinity for various NTRs, can be explored in a high-throughput manner.

The K_D_ values we measured for Twin-Strep-tag-mTFP binding to trivalent Strep-Tactin-SpyCatcher, and Strep-tag II-mTFP binding to monovalent Strep-Tactin-SpyCatcher differ from previously reported measurements of either Strep-tag II or Twin-Strep-tag binding to tetravalent Strep-Tactin (Fig. 2a, Supplementary Fig. 5) ^9,11^. The relatively high K_D_ value we measured for Strep-tag II binding to monovalent Strep-Tactin-SpyCatcherΔ is however in line with our observation that activation of Siglec-14 with N-terminal Strep-tag II requires very high surface densities of monovalent Strep-Tactin generic ligand (mean EC_50_ of 1,100,000 generic ligands per cell, Fig. 5d).

The role of receptor clustering in receptor triggering can also be explored by varying the valency of the generic ligand (Fig. 5c). This can be achieved by refolding Strep-Tag II binding and non-binding (dead) Strep-Tactin subunits in varying ratios. Large-scale receptor clustering could also be induced using larger multivalent ligand complexes^40^.

The high density of generic ligand that can be achieved provides an opportunity to use more than one soluble SpyCatcher fusion protein to provide combinations of more than one ligand. For example, a “second generic ligand” could engage co-stimulatory and/or co-inhibitory NTRs or other immune receptors. Such an approach greatly extends our ability to explore not just receptor activation but also signal integration between multiple receptors.

Recently a split receptor system has been described based on expression of a universal cell surface signalling component to which soluble antigen receptors of diverse specificities can be coupled^7^. While such a system is useful for flexible, programmable targeting of effector cells, because the surface density of ligands cannot be precisely varied it is less helpful for studying the mechanisms of immune recognition and receptor triggering.

The generic ligand system, and future work building upon foundations laid here, will aid investigations into the processes that regulate immune signalling and signal integration. In principle, it can also be adapted to analyse the vast array of other receptor-cell surface ligand interactions that exist in a systematic, high-throughput manner.

## Materials and Methods

All data were fitted using GraphPad Prism or FlowJo software. In all cases where sample values were below the machine detection limit, those samples were given a value of 0.

### Constructs

All sequences were verified by Sanger sequencing (Source BioScience or Eurofins Scientific).

#### Strep-tag II/Twin-Strep-tag

Sequences encoding the Igκ leader sequence, Strep-tag II or Twin-Strep-tag with linker (SA<u>WSHPQFEK</u> or SA<u>WSHPQFEK</u>GGGSGGGSGGSA<u>WSHPQFEK</u>), a multiple cloning site, FLAG tag and 3’ stop codon were inserted into the pHR-SIN-BX-IRES-EmGFP lentiviral vector (a gift from Vincenzo Cerundolo, University of Oxford) using 5’ BamHI and 3’ NotI restriction sites and the GENEWIZ Gene synthesis service. This insertion destroyed the plasmid BamHI site and removed the Internal Ribosome Entry Site-Emerald Green Fluorescent Protein (IRES-EmGFP) sequence: pHR-SIN-BX-Strep-tag-II or pHR-SIN-BX-Twin-Strep-tag respectively.

#### Receptors

DNA encoding human SIRPβI (amino acids 30-398), Siglec-14 (amino acids 17-396) or NKp30 (amino acids 19-201) with either 5’ BamHI or XhoI and 3’ BsiWI restriction sites were inserted into the multiple cloning site of pHR-SIN-BX-Twin-Strep-tag. DNA encoding Siglec-14 (amino acids 17-396) was likewise inserted into the multiple cloning site of pHR-SIN-BX-Strep-tag II. DNA encoding human IG4 TCR α and β chains flanking a viral 2A peptide sequence was inserted into the multiple cloning site of pHR-SIN-BX-Twin-Strep-tag such that the tag is at the N terminus of the TCRβ chain. IG4 TCR was amplified without the native β chain signal peptide. The α and β chains also have a C-terminal FLAG tag and HA tag respectively.

#### Adaptor proteins

DNA encoding human DAP12 or FcRγ followed by C-terminal Myc tag was inserted into pHR-SIN-BX-IRES-EmGFP using 5’ BamHI and 3’ XhoI sites, retaining the plasmid IRES-EmGFP sequence.

#### Ligand anchor

Sequence:

**METDTLLLWVLLLWVPGSTGD***YPYDVPDYA*TGGS<u>AHIVMVDAYKPTK</u>GGSGGS *<u>HVSEDFTWEKPPEDPPDSKNTLVLFGAGFGAVITVVVIVVIIKCFCKHRSCFRRNEA SRETNNSLTFGPEEALAEQTVFL</u>*

DNA encoding the Igκ leader sequence (highlighted in bold), HA tag (in italics), SpyTag (underlined), and the extracellular hinge-like region, transmembrane and intracellular regions of mouse CD80 (amino acids 227-306) (in italics and underlined) was inserted into pEE14 using 5’ HindIII and 3’ XbaI restriction sites: pEE14-ligand anchor.

#### HLA-A*02 single chain dimer

##### Sequence

*MSRSVALAVLAILSLSGLEAIQRTPKIQVYSRHPAENGKSNFLNCYV SGFHPSDIEVDLLKNGERIEKVEHSDLSFSKDWSFYLLYYTEFTPTEKDEYACRV NHVTLSQPKIVKWDRDM***GGGGSGGGGSGGGGS** <u>GSHSMRYFFTSVSRPGRGEPRFIAVGYVDDTQFVRFDSDAASQRM EPRAPWIEQEGPEYWDGETRKVKAHSQTHRVDLGTLRGYYNQSEAGSHTVQRMYG CDVGSDWRFLRGYHQYAYDGKDYIALKEDLRSWTAADMAAQTTKHKWEAAHVAEQ LRAYLEGTCVEWLRRYLENGKETLQRTDAPKTHMTHHAVSDHEATLRCWALSFYP AEITLTWQRDGEDQTQDTELVETRPAGDGTFQKWAAVVVPSGQEQRYTCHVQHEG LPKPLTLRWEPGSQPTIPIVGIIAGLVLFGAVITGAVVAAVMWRRKSSDRKGGSY SQAASSDSAQGSDVSLTACKV</u>

DNA encoding human β-2 microglobulin (in italics) and HLA-A*02 (amino acids 25-365) (underlined) separated by a GS linker ([GGGGS]_3_) (in bold) was inserted into pDisplay using 5’ HindIII and 3’ XhoI restriction sites to create a single chain dimer (SCD) : pDISPLAY-SCD.

#### Strep-Tactin, dead streptavidin-SpyCatcher and variants

##### Strep-Tactin sequence

MAEAGITGTWYNQLGSTFIVTAGADGALTGTYVTARGNAESRYVLTGRYDSAPATDG SGTALGWTVAWKNNYRNAHSATTWSGQYVGGAEARINTQWLLTSGTTEANAWKSTLV GHDTFTKVKPSAAS

##### Dead streptavidin-SpyCatcher sequence

Dead streptavidin is underlined, SpyCatcher is in italics, and the polyaspartate sequence is in bold.

<u>MAEAGITGTWYAQLGDTFIVTAGADGALTGTYEAAVG</u>**DDDGDDDGDDDG** <u>AESRYVLTGRYDSAPATDGSGTALGWTVAWKNNYRNAHSATTWSGQYVGG AEARINTQWLLTSGTTEANAWKSTLVGHDTFTKVKPSAAS</u> GSGSG*DYDIPTTENLYFQGAMVDTLSGLSSEQGQSGDMTIEEDSAT HIKFSKRDEDGKELAGATMELRDSSGKTISTWISDGQVKDFYL YPGKYTFVETAAPDGYEVATAITFTVNEQGQVTVNGKATKGDAHI*

##### Dead streptavidin sequence

MAEAGITGTWYAQLGDTFIVTAGADGALTGTYEAAVGNAESRYVLTGRYDSAPATDG SGTALGWTVAWKNNYRNAHSATTWSGQYVGGAEARINTQWLLTSGTTEANAWKSTLV GHDTFTKVKPSAAS

##### Strep-Tactin-SpyCatcherΔ sequence

Strep-Tactin is underlined, SpyCatcherΔ is in italics and the polyaspartate sequence is in bold.

<u>MAEAGITGTWYNQLGSTFIVTAGADGALTGTYVTARGNAESRYVLTGRYD SAPATDGSGTALGWTVAWKNNYRNAHSATTWSGQYVGGAEARINTQWLLT SGTTEANAWKSTLVGHDTFTKVKPSAAS</u> **DDDGDDDGDDDD** *SATHIKFSKRDEDGKELAGATMELRDSSGKTISTWISDGQVKDFYLYPGKYTF VETAAPDGYEVATAITFTVNEQGQVTVNGKATKGDAHI*

pET21-Strep-Tactin, pET21-dead streptavidin-SpyCatcher, and pET21-dead streptavidin (Addgene plasmid #20859) were gifts from Mark Howarth, University of Oxford^17^.

The dead streptavidin sequence of pET21-dead streptavidin-SpyCatcher matches Addgene plasmid #59547 with polyaspartate sequence in the streptavidin 3/4 loop for anion exchange chromatography^40^. The C-terminal SpyCatcher sequence is different however, as per Addgene plasmid #35044^12^. pET21-Strep-Tactin-SpyCatcherΔ was created by amplification of Strep-Tactin and truncated SpyCatcher (as in Addgene plasmid #59547) sequences and insertion either side of a polyaspartate sequence^40^. SpyCatcher truncation does not significantly affect SpyTag-SpyCatcher reaction efficiency^16^.

### Cell lines

#### THP-1, Jurkat, HEK293T cell lines

THP-1, Jurkat reporter (enhanced Green Fluorescent Protein (eGFP) production under the control of the NFκB promoter), and HEK293T cells were maintained in RPMI-1640 (Sigma-Aldrich # R8758) media supplemented with 10% foetal bovine serum (FBS) and 100 U mL^*-*1^ penicillin/streptomycin (Thermo Fisher Scientific # 15140122) at 37 °C in a 5% CO_2_-containing incubator. Jurkat NFκB-driven eGFP reporter and Jurkat NFκB-driven eGFP reporter IG4 TCRα/β CD8α/β cells were a gift from Peter Steinberger and Wolfgang Paster, Medical University of Vienna^26,27^.

#### CHO cell lines

Chinese Hamster Ovary (CHO) mock cells were maintained in DMEM (Sigma-Aldrich # D6429) supplemented with 5% FBS and 100 U mL^*-*1^ penicillin/streptomycin. CHO ligand anchor cells were maintained in L-Glutamine-free DMEM (Sigma-Aldrich # D6546) supplemented with 5% dialysed FBS (dialysed thrice against 10 L PBS), 100 U mL^*-*1^ penicillin/streptomycin, 1x GSEM supplement (Sigma-Aldrich # G9785) and 50 µM L-Methionine sulfoximine (Sigma-Aldrich # M5379). CHO ligand anchor HLA-A*02 cells were maintained in the above media additionally supplemented with 1 mg mL^*-*1^ G-418 (Thermo Fisher Scientific # 10131027). All CHO cell lines were maintained at 37 °C in a 10% CO_2_-containing incubator.

#### Lentiviral transduction of THP-1 and Jurkat cells

Either receptor-expressing lentivector alone, or with the appropriate adaptor-expressing lentivector, was co-transfected with the lentiviral packaging plasmids pRSV-Rev (Addgene plasmd # 12253), pMDLg/pRRE (Addgene plasmd # 12251) and pMD2.g (Addgene plasmd # 12259) into HEK293T cells using X-tremeGENE™ 9 (Roche) as per the manufacturer’s instructions^41^. Lentiviral packaging plasmids were a gift from Didier Trono, École polytechnique fédérale de Lausanne. Two days after transfection, viral supernatants were harvested, filtered and used for the transduction of either THP-1 or Jurkat cells in the presence of 5 µg mL^*-*1^ Polybrene. For CD8α/β-expressing Jurkat cells, Jurkat reporter cells expressing IG4 TCRα/β tagged with Strep-tag II or Twin-Strep-tag were lentivirally transduced as above with pHR-SIN-BX-CD8b-T2A-CD8a (a kind gift of Peter Steinberger and Wolfgang Paster, Medical University of Vienna).

#### Analysing receptor, adaptor, and co-receptor expression using flow cytometry and cell sorting by Fluorescence-Activated Cell Sorting (FACS)

Representative gating strategies for flow cytometry analyses of all cell lines are shown in Supplementary Fig. 7. For all flow cytometry analyses, between 13,500 and 30,000 gated events were analysed.

Cells were analysed for receptor surface expression by flow cytometry using anti-Strep-tag II antibody Oyster 645 (IBA Lifesciences # 2-1555-050), or anti-Strep-tag II antibody (IBA Lifesciences # 2-1507-001) and anti-mouse IgG1 antibody Alexa Fluor 647 (Thermo Fisher Scientific # A-21240), (BD FACSCalibur™, 640 nm laser, FL4 - 661/16 band-pass filter, BD Biosciences). Introduced adaptor expression was tested via expression of EmGFP encoded on the adaptor lentivector (BD FACSCalibur™, 488 nm laser, FL1 - 530/30 band-pass filter, BD Biosciences). IG4 TCRβ and CD8α expression was analysed using anti-TCR Vβ 13.1 antibody FITC (H131; Thermo Fisher Scientific) and anti-CD8α antibody PE (HIT8a; Biolegend) respectively (BD FACSCalibur™, 488 nm laser, FL1 - 530/30 band-pass filter and FL2 - 585/42 band-pass filter, BD Biosciences). Cells were sorted for high expression of receptor ± introduced adaptor or CD8 co-receptor by fluorescence-activated cell sorting (FACS) (MoFlo Astrios, Beckman Coulter).

#### pMHC tetramer staining of Jurkat cells

HLA-A*02:01 heavy chain (residues 1-278) with C-terminal BirA tag and β2-microglobulin were expressed in *Escherichia coli* as inclusion bodies, refolded in the presence of peptide (SLLMWITQV), and purified using size-exclusion chromatography^42^. NY-ESO-1 (157-165; SLLMWITQV) peptide was purchased at >95% purity (GenScript). Purified peptide-HLA-A*02 was biotinylated in vitro by BirA enzyme (Avidity) and mixed with Streptavidin:RPE (Bio-Rad #STAR4A) to create tetramers. Cells were incubated with a below-saturating concentration of pMHC tetramers:RPE and analysed by flow cytometry (BD FACSCalibur™, 488 nm laser, FL2 - 585/42 band-pass filter, BD Biosciences).

#### Transfection of CHO cells with ligand anchor and HLA-A*02

CHO cells were transfected with either pEE14 (CHO mock) or pEE14-ligand anchor (CHO ligand anchor) using Xtreme-GENE 9™ as per the manufacturer’s instructions. A monoclonal population of CHO ligand anchor cells, created by limiting dilution, were transfected with pDISPLAY-SCD (CHO ligand anchor HLA-A*02) using Xtreme-GENE 9™ as per the manufacturer’s instructions. Transfected lines were cultured in the appropriate selection media after 48 hours.

#### Checking ligand anchor and HLA-A*02 expression by flow cytometry and cell sorting by FACS

Cells were analysed for ligand anchor or HLA-A*02 surface expression by flow cytometry using anti-HA-Tag antibody Alexa Fluor 647 (6E2; Cell Signalling Technology) and anti-HLA-A2 FITC antibody (BB7.2; Santa Cruz Biotechnology) respectively (BD FACSCalibur™, 640 nm laser, FL4 - 661/16 band-pass filter, 488 nm laser, FL1 - 530/30 band-pass filter, BD Biosciences). CHO ligand anchor HLA-A*02 cells were sorted for high expression of ligand anchor and HLA-A*02 using FACS (MoFlo Astrios, Beckman Coulter).

### Expression and purification of trivalent Strep-Tactin-SpyCatcher and monovalent Strep-Tactin-SpyCatcherΔ

Individual subunits (Strep-Tactin and dead streptavidin-SpyCatcher or dead streptavidin and Strep-Tactin-SpyCatcherΔ) were expressed in *Escherichia coli* BL21-CodonPlus (DE3)-RIPL cells (Agilent Technologies # 230280) and refolded from inclusion bodies using a modified version of the protocol previously described by Howarth and Ting^43^. Inclusion bodies were washed in BugBuster (Merck Millipore # 70921) supplemented with lysozyme, protease inhibitors, DNase I and magnesium sulphate as per the manufacturers’ instructions. Subunits were then mixed at a 3:1 molar ratio in order to bias refold towards the desired tetramer: Strep-Tactin and dead streptavidin-SpyCatcher to yield trivalent Strep-Tactin-SpyCatcher, or dead streptavidin and Strep-Tactin-SpyCatcherΔ to yield monovalent Strep-Tactin-SpyCatcherΔ. Tetramers were refolded by rapid dilution and precipitated using ammonium sulphate precipitation. Precipitated protein was resuspended in 20 mM Tris pH 8.0, filtered (0.22 µm filter), and loaded onto a Mono Q HR 5/5 column (GE Healthcare Life Sciences). Desired tetramers were eluted using a linear gradient of 0-0.5 M NaCl in 20 mM Tris pH 8.0, concentrated, and buffer exchanged into 20 mM MES, 140 mM NaCl pH 6.0.

### SDS-PAGE and western blotting

Boiled samples were subjected to SDS-PAGE under reducing conditions on NuPAGE 4-12% Bis-Tris protein gels (Thermo Fisher Scientific # NP0322) and stained using Instant*Blue* protein stain (Expedeon).

For western blotting, CHO ligand anchor cells ± trivalent Strep-Tactin-SpyCatcher were lysed in Tris-buffered saline 1% NP40 at 4 °C. Cleared and boiled cell lysates were then subjected to SDS-PAGE under reducing conditions on NuPAGE 4-12% Bis-Tris protein gels. Proteins were transferred to 0.2 µm PVDF membrane (GE Healthcare # 10600022) using a semi-dry blotting system. The membrane was probed with anti-HA tag antibody (6E2; Cell Signalling Technology) followed by IRDye 680RD goat anti-mouse IgG (LI-COR Biosciences # 925-68070), and anti-streptavidin antibody (Abcam # Ab6676) followed by anti-rabbit IgG Dylight 800 (Thermo Fisher Scientific # SA5-10036).

Gels and blots were imaged using a LI-COR Odyssey Sa imaging system (LI-COR Biosciences) and images analysed using ImageJ software.

### Biotin-4-fluorescein quenching assay

Valency of trivalent and monovalent Strep-Tactin-SpyCatcher was confirmed using the quenching activity of biotin-4-fluorescein (Sigma-Aldrich #B9431) when bound to Strep-Tactin^44,45^. Trivalent or monovalent Strep-Tactin-SpyCatcher was incubated with a titration of biotin-4-fluorescein concentrations in black, opaque 96 well plates for 30 minutes at 25 °C in PBS 1% BSA. Fluorescence was measured (λ_ex_ 485 nm, λ_em_ 520 nm) using a SpectraMax M5 plate reader (Molecular Devices). Fluorescence values were corrected for background fluorescence before analysis.

Negative control (buffer alone) data were fitted with the linear regression formula (equation 1) where Y is fluorescence (arbitrary units (AU)), M is the gradient, X is the concentration of biotin-4-fluorescein (M) and B is the Y-intercept.

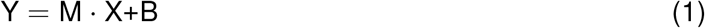

Sample data were fitted with segmental linear regression equation set (equations 2-5) in which X is the biotin-4-fluorescein concentration (M), Y is fluorescence (AU), X0 is the biotin-4-fluorescein concentration at which the line segments intersect (M), slope1 is the gradient of the first line segment, slope2 is the gradient of the second line segment and intercept1 is the Y value at which the first line segment intersects the Y axis. Slope2 was constrained to the value of M calculated for the appropriate negative control (see equation 1).

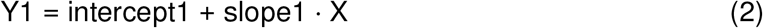

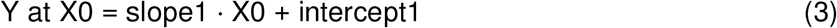

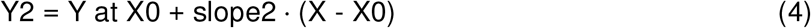

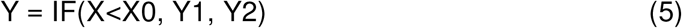

Term X0 was converted into an estimate of number of biotin-binding sites per tetramer using the concentration of Strep-Tactin-SpyCatcher added. For this, we assume complete binding of biotin-4-fluorescein to protein.

### Surface plasmon resonance

Monomeric Teal Fluorescent Protein (mTFP) with C-terminal His tag (6 x His) and either an N-terminal Strep-tag II or Twin-Strep-tag was expressed in *Escherichia coli* and purified by Nickel affinity chromatography^46^.

Affinity measurements were made using a Biacore T200 or 3000 (GE Healthcare). All experiments were performed at 37 °C using a flow rate of 10 µL min^*-*1^ in HBS-EP buffer (0.01 M HEPES buffer (pH 7.4), 0.15 M NaCl, 3 mM EDTA, 0.005% Surfactant P20). SpyTag-containing peptide (<u>AHIVMVDAYKPTK</u>GGSGGSHHHHHHHHHHHH) (SpyTag is underlined), purchased at 95% purity (Peptide Protein Research Ltd.), was immobilised to a sensor chip CM5 (GE Healthcare). Either trivalent Strep-Tactin-SpyCatcher or monovalent Strep-Tactin-SpyCatcherΔ was immobilised to the chip via the SpyTag peptide at various levels. Equilibrium binding was measured for graded concentrations of either Strep-tag II-mTFP or Twin-Strep-tag-mTFP. The K_D_ (M) values were obtained by simultaneously fitting all the data for Twin-Strep-tag-mTFP binding trivalent Strep-Tactin-SpyCatcher (or all the data for Strep-tag II-mTFP binding monovalent Strep-Tactin-SpyCatcherΔ) with equation (6) and constraining the fitting such that the K_D_ value is shared between repeats. Y is the specific binding of injected mTFP fusion protein (Response Units (RU)), Bmax is the maximum specific binding (RU), X is the concentration of injected mTFP fusion protein (M) and h is the Hill slope.

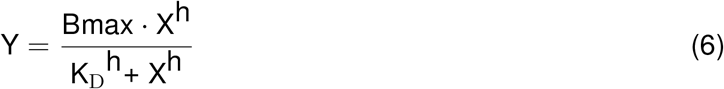

### Generating CHO generic ligand cells

CHO ligand anchor or CHO ligand anchor HLA-A*02 cells (or CHO mock cells as a control) were incubated with various concentrations of trivalent or monovalent Strep-Tactin-SpyCatcher in 20 mM MES, 140 mM NaCl pH 6.0 1% BSA for 10 minutes at 25 °C unless otherwise stated. Unbound Strep-Tactin-SpyCatcher was removed by washing thrice with PBS 1% BSA.

### Measuring generic ligand numbers per cell

To measure the number of generic ligands on a cell population pre-incubated with a single concentration of trivalent or monovalent Strep-Tactin-SpyCatcher the above biotin-4-fluorescein fluorescence quenching assay was used. A titration of biotin-4-fluorescein was incubated with known numbers of cells pre-incubated with trivalent or monovalent Strep-Tactin-SpyCatcher (or buffer alone as a control). The X0 term (calculated from the curve fit using equations 2-5) was converted to an estimate of average generic ligand number per cell using equation (7) where L is the average number of ligands per cell, X0 is the saturation concentration of biotin-4-fluorescein extracted (M), V is the sample volume (L), N_A_ is Avogadro’s constant, C is the number of cells in the sample, and B is the number of biotin-binding sites per ligand. In the case of trivalent Strep-Tactin-SpyCatcher and monovalent Strep-Tactin-SpyCatcherΔ, B = 3 or 1 respectively.

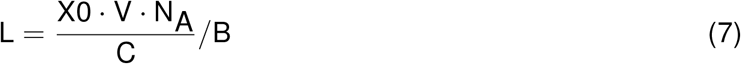

To indicate relative levels of generic ligand per cell, CHO cells pre-incubated with trivalent Strep-Tactin-SpyCatcher or buffer alone were incubated with 2 µM ATTO 647 biotin (ATTO Technology # AD 647-71) pre-mixed with 6 µM biotin (Sigma-Aldrich # B4501) for 30 minutes at 25 °C. The presence of biotin minimises the self-quenching activity of ATTO dye biotin conjugates that we observed (data not shown). When monovalent Strep-Tactin-SpyCatcherΔ was used, cells were incubated with 2 µM ATTO 488 biotin (ATTO Technology # AD 488-71) alone and treated as above. Cells were analysed by flow cytometry (BD FACSCalibur™, 640 nm laser, FL4 - 661/16 band-pass filter, 488 nm laser, FL1 - 530/30 band-pass filter, BD Biosciences). Background median fluorescence intensities (MFIs), or geometric mean fluorescence intensities (gMFIs) when using monovalent Strep-Tactin, where cells were incubated with buffer alone instead of Strep-Tactin-SpyCatcher, were subtracted from all corresponding sample MFI/gMFI values respectively. These values were then fitted with equation (8) where Y is the MFI or gMFI (AU), Bmax is the maximum specific trivalent or monovalent Strep-Tactin-SpyCatcher binding indicated by MFI or gMFI respectively (in AU), X is the concentration of trivalent or monovalent Strep-Tactin-SpyCatcher added (M), and K_D_ we use as the concentration of Strep-Tactin-SpyCatcher that yields 50% maximal binding to CHO cells (M). Whether the median or geometric mean was extracted from flow cytometry analyses is indicated on each graph.

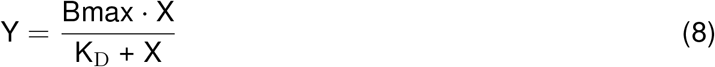

To convert Y values into estimates of average generic ligand number per cell, the ligand number per cell at saturating trivalent or monovalent Strep-Tactin-SpyCatcher concentration calculated from equation (7) was substituted into equation (8) as Bmax. Y values were then re-calculated following this adjustment. Average ligand density (molecules/µm^2^) was calculated from these estimates by assuming a CHO cell surface area of 700 µm^2^. The latter is based on a diameter of 15 µm and an assumed spherical shape^47,48^.

We have independently verified the numbers of generic ligands per cell quantified via this method by flow cytometry using antibodies and IgG quantitation beads (data not shown).

### Down-regulation of generic ligand over time

CHO ligand anchor cells were incubated with 15 µM trivalent Strep-Tactin-SpyCatcher (or buffer alone) as described above and incubated at 37 °C as for stimulation assays for the time points indicated. Cells were analysed for generic ligand surface expression using ATTO 647 biotin as above and normalised to the median fluorescence intensity value at time 0. To calculate the decay, the MFI values were fitted with equation 9 where Y0 is the Y value when X = 0, Plateau is the Y value at which the curve reaches a plateau, X is time in minutes, and K is the rate constant in inverse minutes.

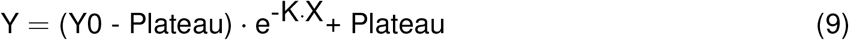

### Measuring number of 9V-HLA-A*02 per cell

The α and β subunits of c58c61 high-affinity IG4 TCR were expressed in *Escherichia coli* as inclusion bodies, refolded in vitro, and purified using size-exclusion chromatography as described previously^31,49^. This protein was then fluorescently labelled with Alexa Fluor 647 NHS Ester (Thermo Fisher Scientific #A37573) and the degree of protein labelling was calculated as per the manufacturer’s instructions. We will refer to this labelled high affinity TCR as IG4hi TCR AF647.

CHO mock or CHO ligand anchor HLA-A*02 cells were incubated with a titration of NY-ESO-1 9V peptide for 1-3 hours at 37 °C as per stimulation assays. Washed cells were incubated with an above saturation concentration of IG4hi TCR AF647 for 1 hour at 4 °C. Cells were analysed by flow cytometry alongside Alexa Fluor 647 fluorescence quantitation beads (Bangs Laboratories #647) (BD FACSCalibur™, 640 nm laser, FL4 - 661/16 band-pass filter, BD Biosciences). Bead MFI values were extracted and used to form a standard curve from which the Jurkat cell specific binding MFIs were interpolated to give estimates of number of 9V-HLA-A*02 per cell, correcting for the degree of labelling of IG4hi TCR AF647. In each experiment the number of 9V-HLA-A*02 per cell for the top two or three cell sub-populations were extrapolated from the standard curve instead of interpolated.

### Cellular functional assays

During all functional assays, samples of prepared CHO cells were used for estimating the number of generic ligands or 9V-HLA-A*02 per cell.

#### THP-1 cellular assays

CHO mock or CHO ligand anchor cells (2 x 10^5^) pre-incubated with trivalent Strep-Tactin-SpyCatcher or buffer alone were mixed with Twin-Strep-tagged receptor- and adaptor-expressing THP-1 or untransduced THP-1 cells (1 x 10^5^) in DMEM 5% FBS, 100 U mL^*-*1^ penicillin/streptomycin, 2 µg mL^*-*1^ avidin. Cells were incubated in a 37 °C 10% CO_2_-containing incubator for 20 hours. Supernatants were harvested and assayed for IL-8 by ELISA (Thermo Fisher Scientific #88-8086-77). Alternatively, CHO ligand anchor cells (2 x 10^5^) pre-incubated with monovalent Strep-Tactin-SpyCatcherΔ or buffer alone were mixed with THP-1 Siglec-14-Strep-tag II DAP12 cells (1 x 10^5^) in DMEM 5% FBS, 100 U mL^*-*1^ penicillin/streptomycin, 2 µg mL^*-*1^ avidin. Cells were then incubated, supernatant harvested, and IL-8 secretion assayed as above.

#### Jurkat cellular assays

CHO mock or CHO ligand anchor HLA-A*02 cells (2 x 10^5^) pre-incubated with trivalent Strep-Tactin-SpyCatcher or buffer alone were mixed with untransduced Jurkats or Jurkat IG4 TCRα/β-Twin-Strep-tag cells (1 x 10^5^) in DMEM 5% FBS, 100 U mL^*-*1^ penicillin/streptomycin, 2 µg mL^*-*1^ avidin. Alternatively, CHO mock or CHO ligand anchor HLA-A*02 cells (2 x 10^5^) were mixed with untransduced Jurkat cells or Jurkat IG4 TCRα/β-Twin-Strep-tag cells (1 x 10^5^), and NY-ESO-1 9V peptide in DMEM 5% FBS, 100 U mL^*-*1^ penicillin/streptomycin, 2 µg mL^*-*1^ avidin. Cells were incubated in a 37 °C 10% CO_2_ incubator for 20 hours. Cells were harvested, incubated with anti-CD45 antibody Alexa Fluor 647 (F10-89-4; Bio-Rad Laboratories) and analysed by flow cytometry (BD FACSCalibur™, 640 nm laser, FL4 - 661/16 band-pass filter, 488 nm laser, FL1 - 530/30 band-pass filter, BD Biosciences). A representative gating strategy for flow cytometry analyses is shown in Supplementary Fig. 7. For flow cytometry analyses post cellular stimulation, at least 9,300 gated events were analysed in each experiment.

#### Analysis

For THP-1 cell assays, IL-8 concentrations in negative controls (where CHO cells were pre-incubated with buffer alone instead of Strep-Tactin-SpyCatcher) were subtracted from corresponding sample IL-8 concentrations to correct for background levels. Dose-response curves were then fitted with equation (10) where Y is the measured cell response (pg mL^*-*1^), Bottom and Top are the minimum and maximum cell response respectively (pg mL^*-*1^), EC_50_ is the ligand number per cell that yields a half maximal response, X is the number of generic ligands per cell, and Hill slope relates to the steepness of the curve.

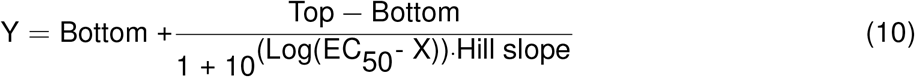

For Jurkat cell assays, only CD45+ cells were analysed for eGFP. Negative control populations were used to set a gate on eGFP expression (see Supplementary Fig. 7). The percentages of CD45+ cells positive for eGFP were extracted and corrected for corresponding background % eGFP+ Jurkat cells (samples where CHO cells were incubated with buffer alone instead of Strep-Tactin or peptide). Dose-response curves were fitted with equation (10) where Y is the measured cell response (% or AU), Bottom and Top are the minimum and maximum cell response respectively (% or AU), EC_50_ is the ligand number per cell that yields a half maximal response, X is the number of generic ligands or 9V-HLA-A*02 per cell, and Hill slope relates to the steepness of the curve. When comparing generic ligand and native ligand dose response curves the Y values were normalised to the individual data set maximal response, giving the maximum a value of 1.

## Acknowledgements

We acknowledge Mark Howarth for providing Strep-Tactin, streptavidin, dead streptavidin, SpyTag and SpyCatcher constructs and for helpful discussions and advice; Peter Steinberger and Wolfgang Paster for providing Jurkat reporter cell lines and Adaptimmune Limited for providing the c58c61 TCR. We thank Mikhail Kutuzov for providing 9V-HLA-A*02 and other members of the Dushek group for providing 9V-HLA-A*02 tetramers. We also thank Marion H Brown and Omer Dushek for helpful discussion.

This work was supported by two Wellcome Trust PhD Studentships (E.M.D., grant reference: 097108/Z/11/Z, R.L.P., grant reference: 099812/Z/12/Z), a Wellcome Trust Senior Investigator Award (P.A.v.d.M., grant reference: 101799/Z/13/Z) and a Nuffield Medical Fellowship from the Australian Academy of Science (J.G., grant reference: #1016848).

## Author Contributions

J.G. and P.A.v.d.M. conceived the study and were responsible for overall design. E.M.D., M.I.B., S.M.B., M.J.B., B.d.W., R.L.P., and J.G. performed research. E.M.D., M.I.B., M.J.B., B.d.W., and J.G. analysed data. E.M.D., P.A.v.d.M., and J.G. wrote the article.

## Supplemental information

**Supplementary Figure 1:**
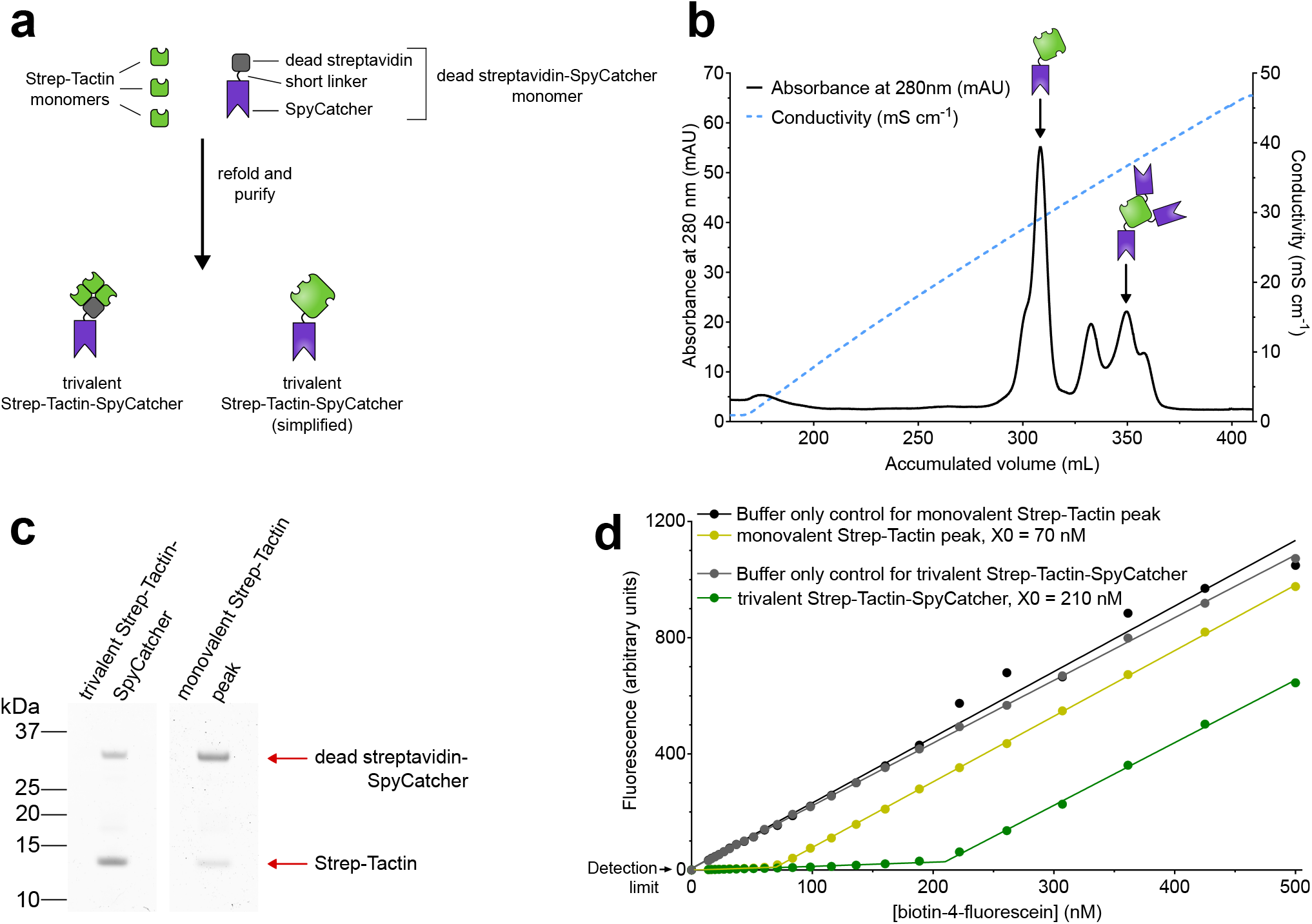
Preparation of trivalent Strep-Tactin-SpyCatcher. (**a**) Trivalent Strep-Tactin-SpyCatcher is synthesised by refolding mixtures of bacterially-produced Strep-Tactin and dead streptavidin-SpyCatcher monomers in a 3:1 ratio. This desired tetramer is shown in schematic form alongside a more simplified cartoon. (**b**) An anion exchange chromatogram showing elution of the predicted trivalent Strep-Tactin-SpyCatcher peak alongside other configurations including a monovalent Strep-Tactin tetramer with a single Strep-tag II binding site and three SpyCatchers. (**c**) SDS-PAGE analysis of the eluted anion exchange chromatography peaks predicted to contain trivalent Strep-Tactin-SpyCatcher and monovalent Strep-Tactin. The samples were boiled prior to loading and the gel was stained with Coomassie to show the relative proportion of subunits. Densitometry was performed on Strep-Tactin and dead streptavidin-SpyCatcher bands, the values normalised for subunit molecular weights, and then converted into a Strep-Tactin : dead streptavidin-SpyCatcher subunit ratio. Trivalent Strep-Tactin-SpyCatcher = 4.7:1 (expected 3:1), monovalent Strep-Tactin peak = 0.6:1 (expected 0.33:1). (**d**) Trivalent Strep-Tactin-SpyCatcher or the predicted monovalent peak for comparison (50 nM) was incubated with a titration of biotin-4-fluorescein in a fluorescence quenching assay. Inflection point X values (X0) are shown.

**Supplementary Figure 2:**
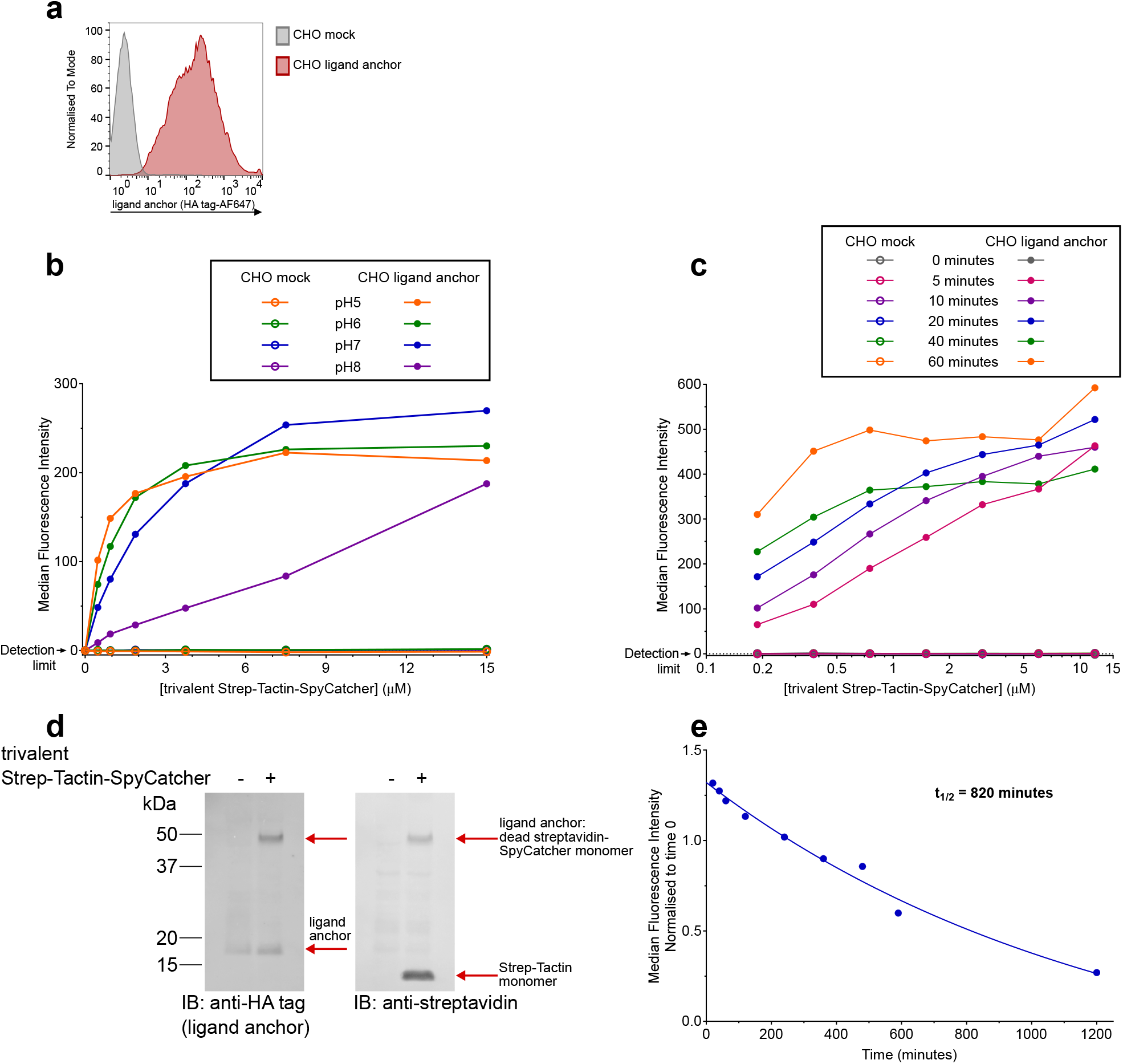
Optimal conditions for generating the complete generic ligand. (**a**) Ligand anchor expression by transfected CHO cells as determined using antibody to N-terminal HA tag and flow cytometry (CHO mock - mock transfected). The efficiency of CHO ligand anchor:trivalent Strep-Tactin-SpyCatcher coupling at 25°C under different pH conditions (**b**) or with varying cell-protein incubation times before washing (**c**) is shown. Cells were incubated with ATTO 647 biotin to indicate generic ligand levels. Median fluorescence intensity values, extracted from flow cytometry analyses, are shown as a function of trivalent Strep-Tactin-SpyCatcher concentration. (**d**) Ligand anchor:trivalent Strep-Tactin-SpyCatcher binding is covalent. Boiled lysates of CHO ligand anchor cells incubated with trivalent Strep-Tactin-SpyCatcher or buffer only were analysed by western blotting. IB, immunoblot. The Strep-Tactin-SpyCatcher tetramer dissociates upon boiling and so the ligand anchor is visualised coupled to dead streptavidin-SpyCatcher subunit only. (**e**) Cell surface down-regulation of the generic ligand over time following reconstitution is visualised using ATTO 647 biotin. Median fluorescence intensities, extracted from flow cytometry analyses, are shown normalised to the MFI at time 0 which was given a value of 1. The mean half-life from two independent experiments (range = 780-860 minutes, n = 2) is shown. The generic ligand cell surface levels appear to rise within the first 20 minutes post-reconstitution, visualised as an increase in MFI. This may reflect a proportion of trivalent Strep-Tactin-SpyCatcher that is in contact with, but not yet covalently bound to, ligand anchor during the initial incubation and so is removed during the process of analysing ligand cell surface levels. Incubating the cells at 37 °C post-reconstitution may allow this proportion of protein to covalently, irreversibly bind to the ligand anchor and thereby lead to an apparent increase in cell surface levels.

**Supplementary Figure 3:**
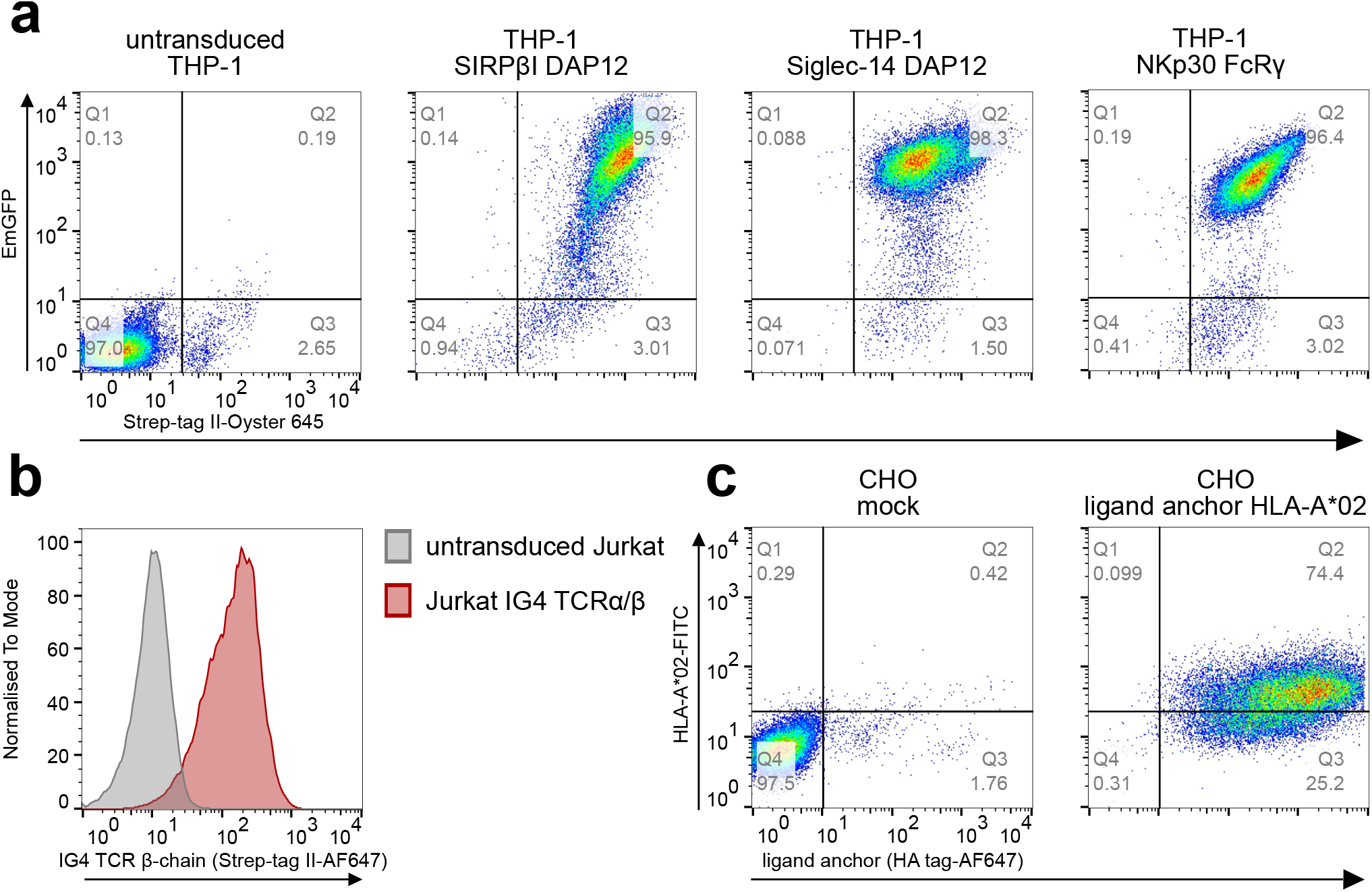
Twin-Strep-tagged receptor and associated adaptor expression and CHO ligand anchor HLA-A*02 expression. (**a**) Expression of one of three Twin-Strep-tagged receptors and appropriate adaptor by THP-1 cells. Using flow cytometry, receptor expression was analysed using anti-Strep-tag II antibody. Expression of the exogenous, introduced adaptor was inferred using an Internal Ribosome Entry Site-Emerald Green Fluorescent Protein (IRES-EmGFP) sequence. (**b**) Expression of IG4 TCRα/β-Twin-Strep-tag by Jurkat cells as shown using anti-Strep-tag II antibody and flow cytometry. (**c**) Expression of the generic ligand anchor and HLA-A*02 single chain dimer by CHO cells shown using anti-HA tag antibody and anti-HLA-A*02 antibody respectively. Numbers indicate percentage of events in each quadrant.

**Supplementary Figure 4:**
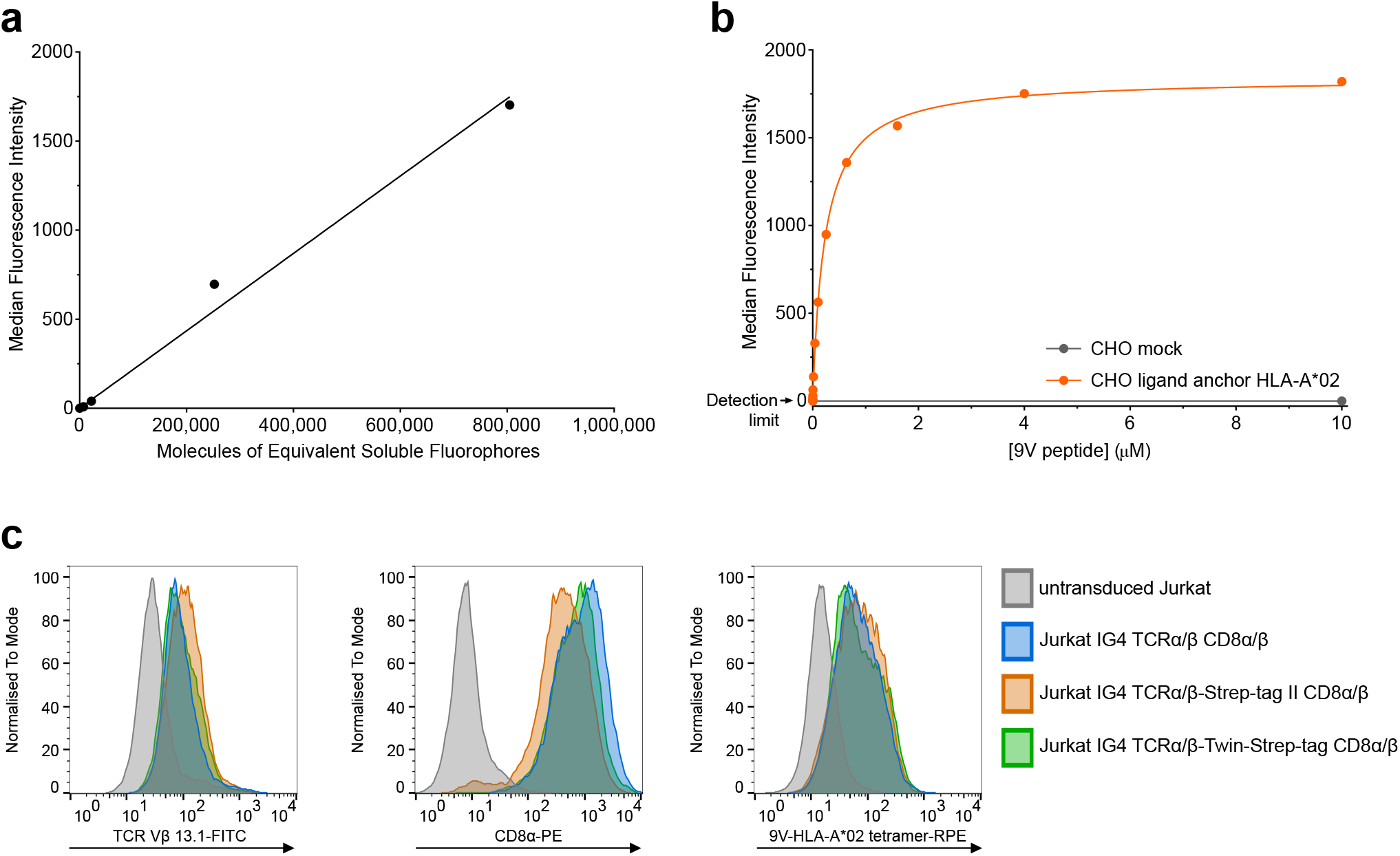
Quantification of 9V-HLA-A*02 per cell and demonstration that Twin-Strep-tag does not interfere with TCR-9V-HLA-A*02 binding. (**a**) Median fluorescence intensity values from flow cytometry analysis of Alexa Fluor 647 fluorescence quantitation beads used to create a standard curve. (**b**) A relative indication of the level of 9V-HLA-A*02 per cell as a function of 9V peptide concentration added to cells. Median fluorescence intensity values extracted from flow cytometry analyses of cells incubated with soluble IG4 high affinity TCRα/β Alexa Fluor 647 are shown. (**c**) Jurkat reporter cells expressing CD8α and β, and IG4 TCRα/β either non-tagged or tagged with Strep-tag II or Twin-Strep-tag show comparable levels of TCRβ chain (left) and CD8α (centre) expression and 9V-HLA-A*02 tetramer binding (right).

**Supplementary Figure 5:**
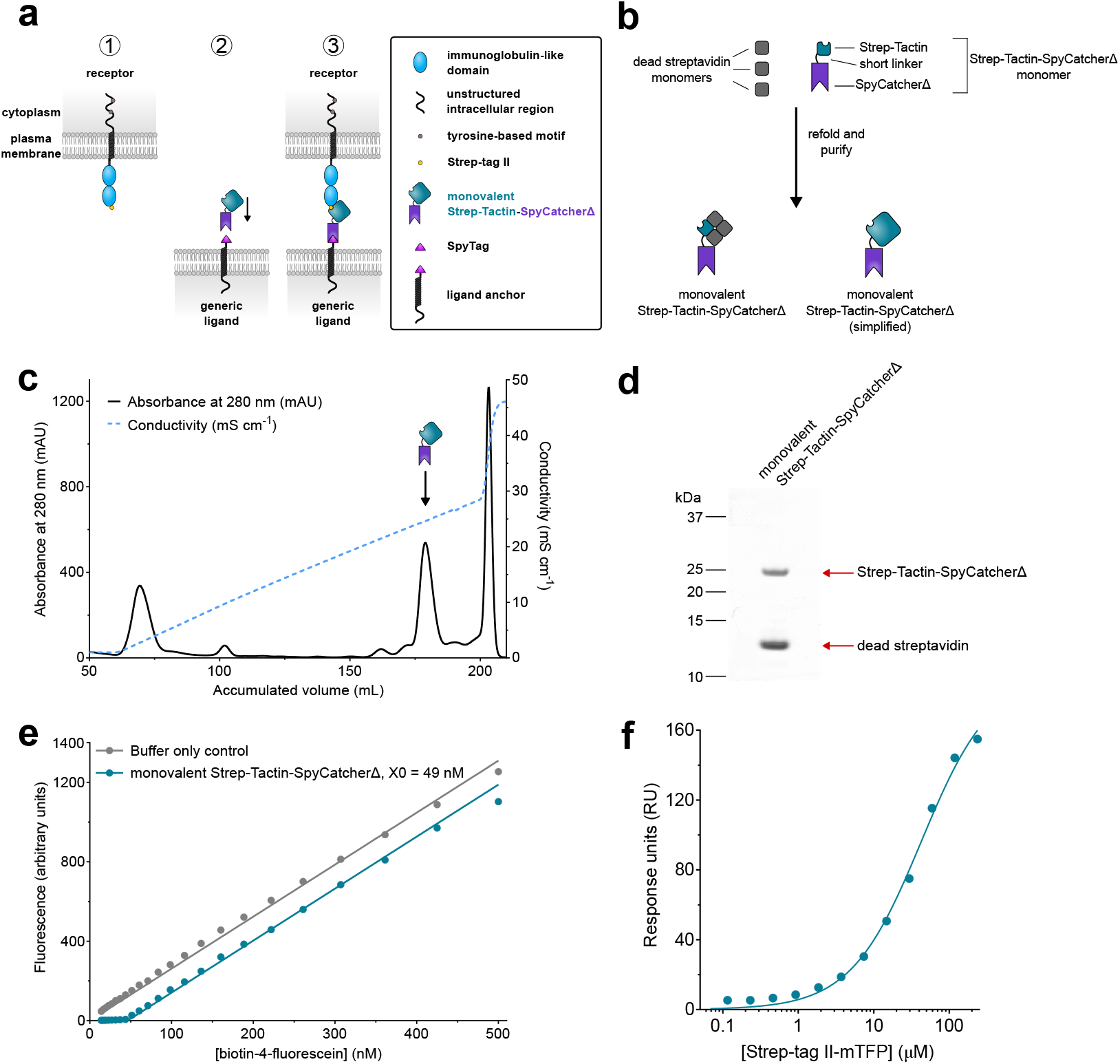
Lowering receptor-generic ligand affinity using Strep-tag II-tagged receptor and monovalent Strep-Tactin-SpyCatcheΔ. (**a**) (1) The receptor is constructed with N-terminal Strep-tag II. (2) Soluble monovalent Strep-Tactin-SpyCatcherΔ protein covalently binds to the generic ligand anchor. (3) The single binding site of monovalent Strep-Tactin-SpyCatcherΔ is available for ligation by the Strep-tag II-tagged receptor. (**b**) Monovalent Strep-Tactin-SpyCatcherΔ is synthesised by refolding mixtures of bacterially-produced Strep-Tactin-SpyCatcherΔ and dead streptavidin monomers in a 1:3 ratio. (**c**) An anion exchange chromatogram showing elution of the predicted monovalent Strep-Tactin-SpyCatcherΔ peak alongside other configurations. (**d**) SDS-PAGE analysis of the eluted anion exchange chromatography peak predicted to contain monovalent Strep-Tactin-SpyCatcherΔ. Densitometry was performed on the bands, the values normalised for subunit molecular weights, and then converted into a Strep-Tactin-SpyCatcherΔ : dead streptavidin subunit ratio (1:3.7, expected 1:3). (**e**) Monovalent Strep-Tactin-SpyCatcherΔ (50 nM) was incubated with a titration of biotin-4-fluorescein in a fluorescence quenching assay. The inflection point (X0) is shown. (**f**) Representative equilibrium binding from surface plasmon resonance of Strep-tag II-mTFP flown over immobilised monovalent Strep-Tactin-SpyCatcherΔ at 37 °C. The K_D_ (SEM) for the collated data from 3 independent experiments (n = 6) is 43 µM (4.5 µM) and the mean Hill slope (SEM) is 0.97 (0.085) to 2 s.f.

**Supplementary Figure 6:**
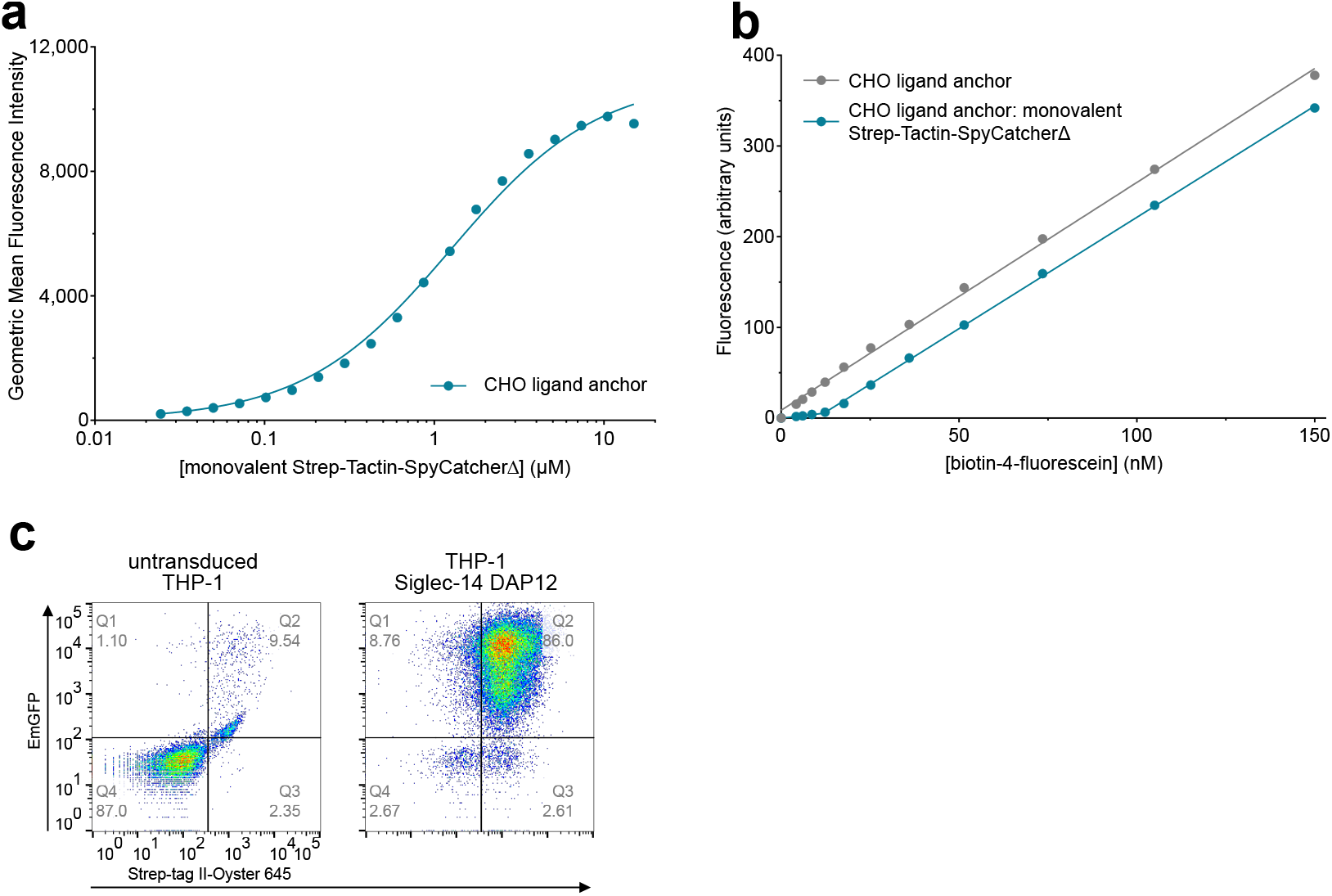
Quantification of monovalent Strep-Tactin-SpyCatcherΔ proteins per CHO ligand cell. (**a**) A relative indication of the level of generic ligand per cell as a function of monovalent Strep-Tactin-SpyCatcherΔ concentration. Geometric mean fluorescence intensity values from flow cytometry analyses of cells incubated with ATTO 488 biotin are shown. (**b**) CHO ligand anchor cells pre-incubated with monovalent Strep-Tactin-SpyCatcherΔ or buffer alone were incubated with a titration of biotin-4-fluorescein in a fluorescence quenching assay. (**c**) Expression of Siglec-14-Strep-tag II and exogenous DAP12 by THP-1 cells using anti-Strep-tag II antibody and an IRES-EmGFP sequence respectively and flow cytometry. Percentages of events in each quadrant are shown.

**Supplementary Figure 7:**
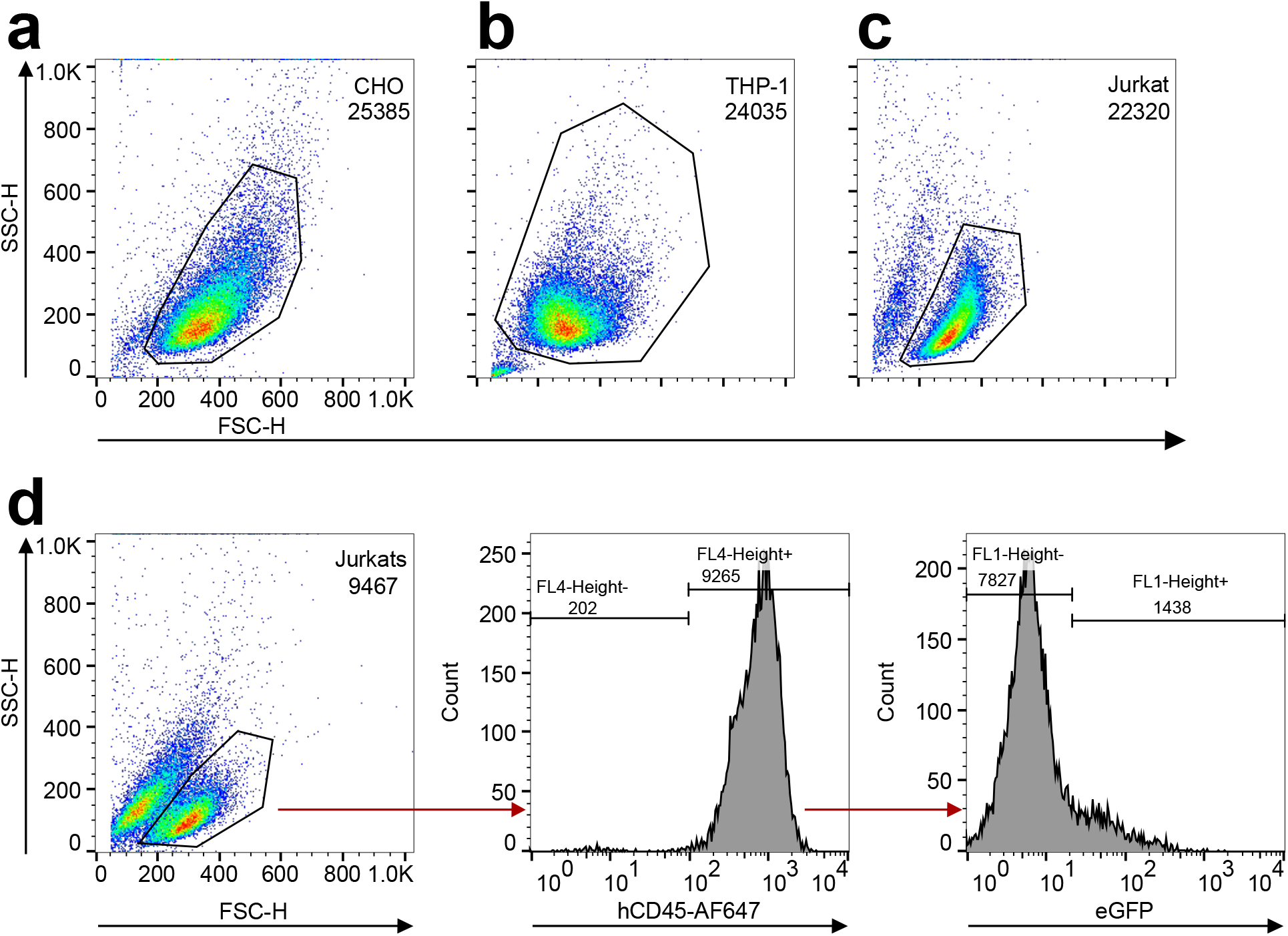
Representative gating strategies. Representative forward and side scatter gating strategies of CHO cells (**a**), THP-1 cells (**b**), and Jurkat cells (**c**). For Jurkat cell stimulation assays all events were gated on forward and side scatter (**d** left), followed by CD45 expression (middle), followed by eGFP expression (right). The number of events in each gate are indicated on each panel.

